# Hidden state inference requires abstract contextual representations in ventral hippocampus

**DOI:** 10.1101/2024.05.17.594673

**Authors:** Karyna Mishchanchuk, Gabrielle Gregoriou, Albert Qü, Alizée Kastler, Quentin Huys, Linda Wilbrecht, Andrew F. MacAskill

## Abstract

The ability to form and utilize subjective, latent contextual representations to influence decision making is a crucial determinant of everyday life. The hippocampus is widely hypothesized to bind together otherwise abstract combinations of stimuli to represent such latent contexts, and to allow their use to support the process of hidden state inference. Yet, direct evidence for this remains limited. Here we show that the CA1 area of the ventral hippocampus is necessary for mice to perform hidden state inference during a 2-armed bandit task. vCA1 neurons robustly differentiate between the two abstract contexts required for this strategy in a manner similar to the differentiation of spatial locations, despite the contexts being formed only from past probabilistic outcomes. These findings offer insight into how latent contextual information is used to optimize decision-making processes, and emphasize a key role of the hippocampus in hidden state inference.

## INTRODUCTION

Animals including humans and rodents have a remarkable capacity for maintaining optimal behavior when faced with a constantly changing environment. This is thought to be achieved by processing otherwise stochastic and volatile information into stable latent representations^1–3^. Once learnt, these subjective representations can be used to infer the most likely context in an otherwise ambiguous situation (often called ‘hidden state inference’), and thus guide optimal behavior^4–10^. Crucially, deficits in hidden state inference are a core feature of many of the most debilitating neural disorders^11–15^. However, while there is much understanding about how directly observable cues or events can be used to guide behavior, the means by which subjective, latent contexts are represented and utilized in the brain remains unclear.

Seminal work has shown that the hippocampus organizes sensory stimuli into representations of discrete spatial locations and contexts to guide flexible behavior^16–18^. This has led to the hypothesis that hippocampal circuitry may play a crucial role in hidden state inference more generally – through the creation and utilization of subjective, often non-spatial latent contexts^1,3,19,20^. However, there is little direct investigation of how the hippocampus represents such information at the neural level, and how this might support behavior.

To address this issue, we used a ‘2-armed bandit’ task in mice - a commonly used probabilistic serial reversal learning task^9,12,21–27^. In this task subjects are presented with a choice of two levers, each with different probabilities of reward, and must use trial and error to identify the best option (the high probability choice). Importantly, after a variable number of correct choices – unsignaled to the participant – the contingencies of the levers switch. Therefore, subjects must both identify and exploit the current high probability lever, but also be able to rapidly switch their choice after a switch in contingencies. Optimal performance in this task can be achieved using hidden state inference – where the participant forms two latent contexts (‘right lever high’ or ‘left lever high’) to guide behavior, based solely on the frequency of past probabilistic outcomes^4,9,12^. We therefore hypothesized that the representation and use of these two latent contexts during the task would be critically dependent on the hippocampus.

In this study we first show that mice use hidden state inference to perform the 2-armed bandit task. We then show that neurons in the ventral CA1 area of the hippocampus are required for both the behavioral use of state inference, and its influence on dopamine dynamics. Finally, we show that this is due to the activity of vCA1 neurons robustly differentiating the two latent contexts that make up the task in a similar manner to the classically observed differentiation of spatial locations.

## RESULTS

### An operant 2-armed bandit task for mice

To provide a framework in which to examine the representation of subjective, latent contexts, we implemented an operant version of a 2-armed bandit task for mice (**Figure 1A**)^9,21–23^. In the full version of the task, mice initiate a trial with a nose poke, and are then presented with two retractable levers either side of the nose port. Pressing one of these levers resulted in reward (6 µl of 10 % sucrose solution) with 70 % probability (high-probability lever), while pressing the opposite lever was rewarded with 10 % probability (low-probability lever). The reward contingencies reversed after 10 to 32 high-probability lever choices.

**Figure 1.**
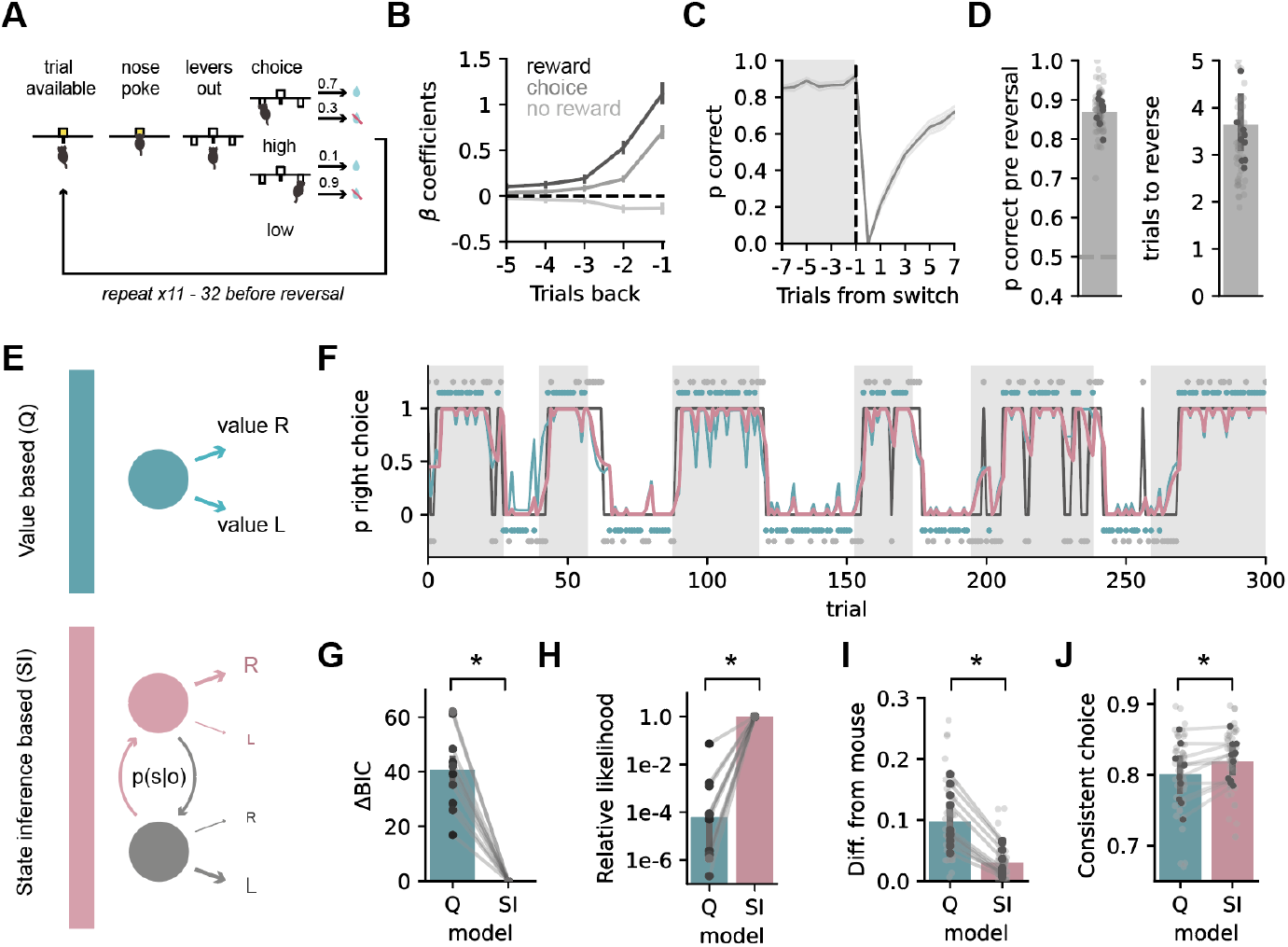
Mice utilize a contextual, state inference strategy to solve a 2-armed bandit task. **A**. Schematic of the task. **B**. Regression coefficients from a logistic regression model predicting animal’s choice on a given trial based on rewarded choice (black), unrewarded choice (light grey) or outcome-independent choice (grey) history. **C**. Average proportion of high probability choices around a switch in contingency. **D**. Left, proportion of high probability choices over the 8 trials before the block reversal. Right, number of trials to switch to the new high probability choice following block reversal. **E**. Schematics of Q and SI model strategies. **F**. Example behavioral session. Grey boxes show blocks of the right lever being the high probability choice; black trace shows animals’ choices (top = right lever press, bottom = left lever press); probability of right choices predicted by the Q (blue) or SI (pink) models are superimposed; blue and grey dots at top and bottom show rewarded and non-rewarded trials. **G**. Summary of model fits to Q and SI models using Bayesian Information Criterion. Note for all mice Q provides a poorer fit. **H**. As in (**G**) but using relative likelihood. **I**. Difference between mouse switching behavior at different trial histories, and that of simulations utilizing either Q or SI strategies. Note for each mouse behavior is closer to SI simulations. **J**. Proportion of choices consistent with a Q or SI strategy. Dark points show individual mice, light points show individual sessions. Error bars represent s.e.m. across animals (n = 10).

Following several stages of training (see Methods), mice reached high performance, and used a combination of past reward and choice to guide their behavior (**Figure 1B,C**). Mice could closely track the identity of the high-probability lever (∼90% correct at the end of each contingency block) and quickly captured the changes in the reward contingencies (mice chose the new lever above chance ∼3 trials after a switch in contingency, **Figure 1D**).

### Mice utilize a state inference strategy to perform a 2-armed bandit task

We next asked what behavioral strategy best described mouse behavior during the task. We focused on a comparison between agents that presume a single context, and incrementally update the value of each choice on a trial-by-trial basis, often termed Q-learning agents (Q); and agents that use state inference (SI), that infer the probability of being in one of two latent contexts, and use this to estimate the optimal choice (**Figure 1E**)^8,9,25,28^. Importantly, we also performed a comprehensive comparison of the most commonly utilized models (**Sup. Figure 1**), which confirmed that Q and SI agents were the strategies that best recapitulated mouse behavior (see Methods).

We found across all sessions and all mice that models utilizing SI consistently best described mouse behavior. We found equivalent results using several complementary metrics. First, we used measurements of the quality of fit such as Bayesian Information criterion (BIC, **Figure 1G**) and relative likelihood (**Figure 1H**). Second, we found that the probability of a mouse switching its choice was strongly shaped by different choices and their outcomes on the past three trials^27^ (**Figure 1B, Sup. Figure 1**), and this pattern of switching behavior was well recapitulated by SI simulations, but not Q simulations (**Figure 1I**). Third, we looked at trial-to-trial model predictions and found that mice were more likely to choose the option predicted by an SI rather than a Q strategy (**Figure 1J**). Finally, as Q and SI agents utilize distinct strategies for updating choices following a switch in contingencies, there was a marked increase in incongruent trials – trials where Q and SI models had opposite predictions – after a switch in contingency (**Sup. Figure 1**). Consistent with our overall model fits – mouse behavior during incongruent trials most frequently followed predictions from the SI model (**Sup. Figure 1**).

### Dopamine dynamics incorporate state inference during task performance

Our results so far suggest that mice utilize state inference to guide behavior in the 2-armed bandit task. Reward prediction errors (RPE) signaled by dopamine release in the nucleus accumbens core (NAc) have been shown to strongly incorporate predictions from state inference, and such signals are increasingly used as a functional readout of the use of state inference strategies^6,7,9,29,30^.

Therefore, we hypothesized that NAc dopamine would contain equivalent signatures of SI during performance of the 2-armed bandit. To investigate this we expressed the extracellular dopamine sensor dLight1.1^31^ in NAc core (**Figure 2A**), and confirmed that there were increases in dLight signal after rewarded outcomes (CS+), and decreases after non rewarded outcomes (CS-) consistent with dopamine signaling (**Figure 2B,C**)^6,7,9,29,30^. We then used two complementary strategies to show that there was strong and consistent influence of SI on NAc dopamine during task performance.

**Figure 2.**
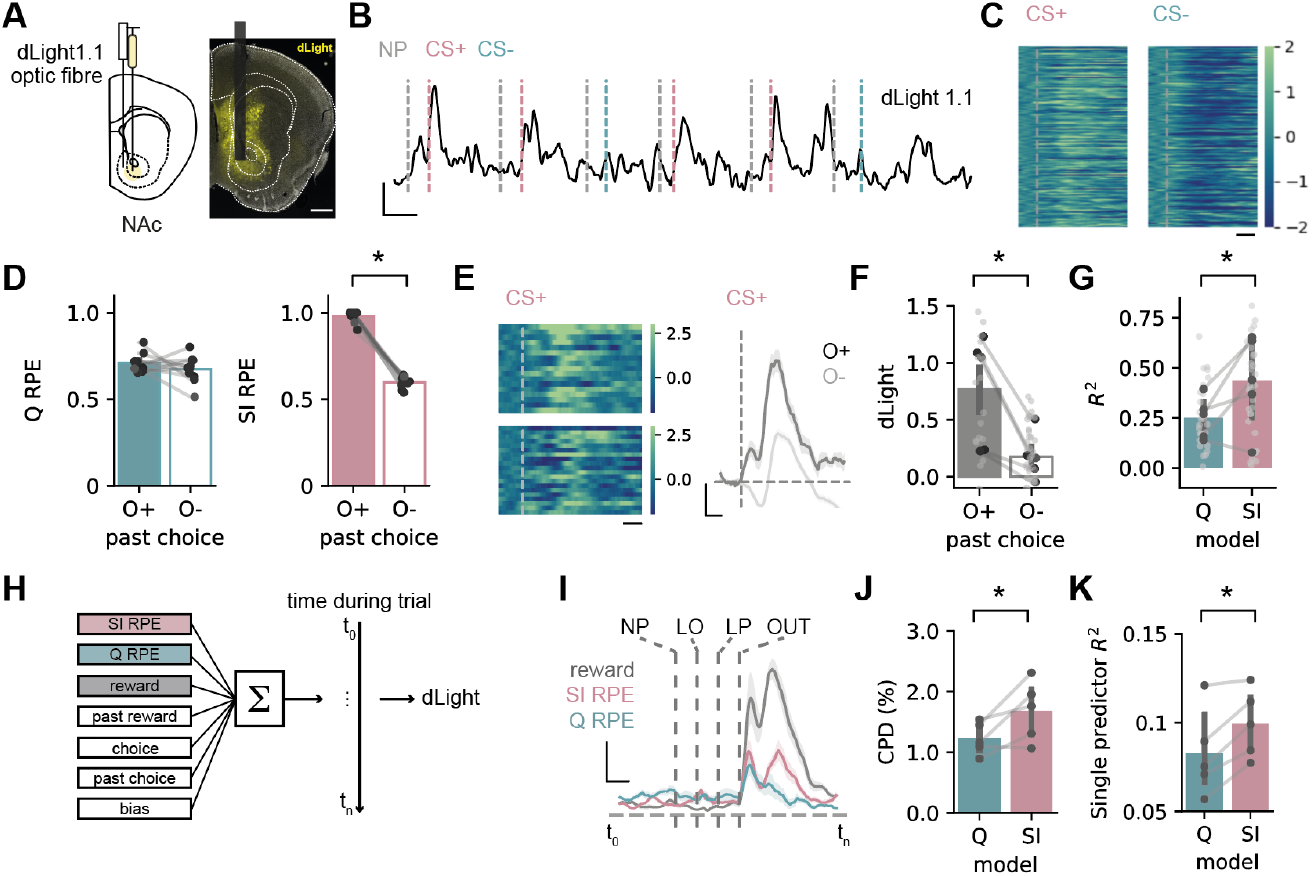
Dopamine in NAc core reflects use of a state inference strategy. **A**. Schematic of dLight1.1 injection and fiber placement in nucleus accumbens core (NAc). Scale bar = 400 μm. **B**. Example recording during task. Grey lines indicate trial initiation, pink lines indicate CS+, and blue lines indicate CS-. Scale bar = 2 Z, 5 seconds. **C**. Z-scored dLight signal time-locked to CS+ or CS-. Scale bar = 1 s. **D**. Prediction error estimates from Q (left) or SI (right) models for trials split according to outcome of past opposite lever press (see text): O+ (reward) or O-(reward omission). Note only SI predictions change. **E**. Left, z-scored dLight signal aligned to CS+ for O+ (top) or O-(bottom) trials. Right, example z-scored dLight signal around CS+ for O+ and O-trials for one session. Scale bar = 1 s (left); 1s, 0.5 zF (right). **F**. Summary of CS+ dLight signal on O+ and O-trials. Note that dLight follows predictions from SI-RPE. **G**. Variance across different past choice and outcome combinations explained by prediction error estimates from either Q or SI models. **H**. Schematic outlining time-lagged ridge regression approach. **I**. Coefficients of partial determination for outcome (grey), Q-RPE (blue) and SI-RPE (pink) at each timepoint. Scale bar = 1 s, 2.5 % **J**. Mean coefficient of partial determination around outcome for Q-RPE and SI-RPE. **K**. Variance explained by Q-RPE or SI-RPE predictors alone. Note consistently higher explained variance from SI-RPE predictor using both metrics. Dark points show individual mice, light points show individual sessions. Error bars represent s.e.m. across animals (n = 5).

First, we took advantage of the fact that Q and SI-based strategies have different mechanics. A Q based agent updates the value of an individual choice only when it is chosen. Therefore, Q-RPE is influenced by a past outcome only if that outcome was associated with the same choice. In contrast, in an SI-based agent the probability of being in a particular latent context is estimated from the past outcome of both choices. Therefore SI-RPE will be influenced by past outcomes from either the same or the opposite choice.

Therefore, we isolated trials where mice chose a different lever than in the previous trial. We then separated these trials dependent on whether the past choice on the opposite lever was rewarded (O+) or not rewarded (O-). We found that consistent with the use of an SI model, dopamine was very different across these two trial histories, and was consistent with estimated RPE obtained from simulations utilizing an SI strategy, but not a Q based strategy (**Figure 2D-F**). Moreover, when we expanded this approach, and compared all possible combinations of trial histories across the same (S) and opposite (O) levers, we found that across all trial types, dopamine was well predicted by SI RPE, and poorly predicted by Q RPE (**Figure 2G, Sup. Figure 2**).

Second, we took a more overarching approach, and used regression to express dLight fluorescence as a sum of responses related to outcome, past outcome, choice and past choice, as well as trial-by-trial estimates of Q-RPE and SI-RPE (**Figure 2H**). Across mice, model weights related to SI-RPE were large, especially around outcome, and consistently explained more variance in dLight signal compared to Q-RPE (**Figure 2I,J**). Moreover, a greater proportion of dLight activity could be explained by SI-RPE alone, when compared to Q-RPE (**Figure 2K**).

Therefore, consistent with our computational modelling, NAc dopamine was strongly influenced by features of an SI strategy. Together, both our behavioral results and dopamine imaging suggest that mice perform inference over 2 abstract task contexts to solve the 2-armed bandit task.

### Ventral CA1 is required for optimal performance in the 2-armed bandit task

Based on our analysis, during performance of the 2-armed bandit task, mice form latent contextual representations that are then used to optimally guide behavior and drive dopamine dynamics. We next used lesions of vCA1 to test our hypothesis that the hippocampus plays a crucial role in utilizing these subjective, non-spatial latent contexts.

We used bilateral expression of caspase under the control of the CaMKii promoter in fully trained mice, to ablate excitatory pyramidal neurons in vCA1 (**Figure 3A**). We found that vCA1 lesions impaired SI-associated behavior, but left behavior guided by other strategies largely intact.

**Figure 3.**
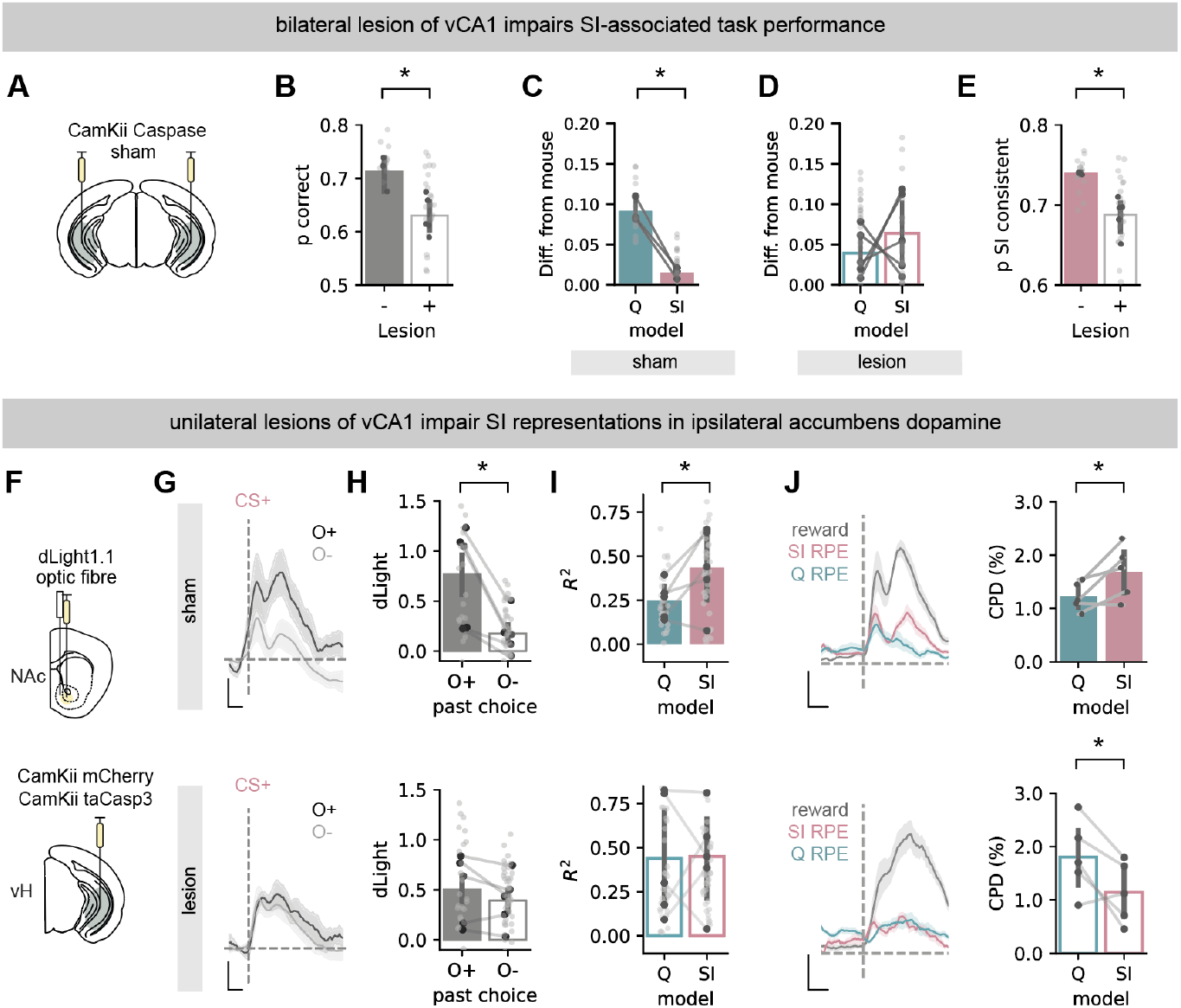
vCA1 is necessary for the neural and behavioral use of state inference. **A**. Schematic of injections to express taCasp3 bilaterally in vCA1 pyramidal neurons. **B**. Summary of high probability choices in sham and caspase lesioned mice. **C**. Difference between mouse switching behavior at different trial histories, and that of simulations utilizing either Q or SI strategies in sham mice. **D**. As in (**C**), but for mice with vCA1 lesions. Note that there is now a marked difference between mouse behavior and SI predictions. **E**. Proportion of choices consistent with SI strategy in sham and lesioned mice. **F**. Schematic of injections to express taCasp3 construct or control fluorophore mCherry in excitatory vCA1 neurons, and dLight1.1 and optic fiber in NAc. **G**. Mean z-scored dLight signal in sham (top) and lesion (bottom) mice, aligned to CS+, and split by different outcomes following opposite lever press in the previous trial (as in Fig 1 K-L). Scale bar = 1s, 0.5 zF. **H**. Summary of CS+ dLight in (**G**) in sham (top) and lesion (bottom) mice. **I**. Variance across different past choice and outcome combinations explained by prediction error estimates from either Q or SI models in sham (top) and lesion (bottom) mice. **J**. Left, coefficients of partial determination for outcome (grey), Q-RPE (blue) and SI-RPE (pink) at each timepoint around cs in sham (top) and lesion (bottom) mice. Right, mean coefficient of partial determination around outcome for Q-RPR and SI-RPE in sham (top) and lesion (bottom) mice. Note consistently higher explained variance from SI-RPE predictor in sham, but this is reversed in lesioned animals. Scale bar = 1 s, 2.5 %. Dark points show individual mice, light points show individual sessions. Error bars represent s.e.m. across animals (n = 5 (sham), n = 5 (lesion)).

When compared to sham injected controls, lesion of vCA1 impaired overall behavior (**Figure 3B**). Similar to our findings in **Figure 1**, we found that the probability of a mouse switching its choice was strongly shaped by different past choices and their outcomes in both sham and lesion groups, particularly in trials with a previous positive outcome (**Sup. Figure 3**). However, while the pattern of switching behavior in sham animals was well recapitulated by SI simulations, and not Q simulations (**Figure 3C**), this was not apparent in lesioned animals (**Figure 3D**), where the ability of an SI strategy to describe switching behavior was markedly reduced. Moreover, when we looked at trial-to-trial model predictions, we found that there was a marked decrease in the proportion of choices consistent with an SI strategy (**Figure 3E**), particularly around switches in contingency (**Sup. Figure 3**), and this change was not apparent when comparing choices consistent with a Q strategy (**Sup. Figure 3**). Together these data suggest lesion of vCA1 results in changes in behavior consistent with a loss of an SI strategy.

### Ventral CA1 is required for the influence of latent context on dopamine dynamics

We next returned to dopamine recordings to gain further insight into the influence of vCA1 lesions on behavioral strategy. If vCA1 was truly shifting behavior due to a loss of the SI representation, we hypothesized that this should be reflected as a loss of SI associated features in NAc dopamine characterized in **Figure 2**^6,7,9,29,30^.

We tested this hypothesis using small, unilateral lesions of excitatory neurons in vCA1 (**Figure 3F**). We performed these lesions unilaterally so as to avoid influence on behavior due to redundancy across hemispheres (**Sup. Figure 3**). Thus, by recording NAc dopamine ipsilaterally to the lesion site, we could monitor the influence of vCA1 on dopamine dynamics, without the confound of altered behavior during our recordings.

Compared to control animals, we found that lesion of vCA1 resulted in NAc dopamine no longer showing features of SI-RPE. This was apparent both by comparing the effect of past trial outcome (**Figure 3G-I**), and also using a regression approach (**Figure 3J**). Therefore, consistent with our hypothesis, in control animals, NAc dopamine contains strong signatures of SI, while lesions of vCA1 result in an almost complete loss of this influence. Together with our behavioral experiments (**Figure 3A-E**), this suggests that vCA1 is required for the use of SI to guide behavior during a 2-armed bandit task.

### vCA1 neurons differentiate choice, expected outcome and latent context during task performance

Our data so far show that i) mice performing a 2-armed bandit task guide their choices by inferring across two latent contexts, defined solely by past choices and their probabilistic outcomes, and ii) that this strategy depends on the activity of vCA1 neurons. During spatial navigation, ‘place cells’ in the CA1 region of the hippocampus fire when an animal occupies a particular spatial location, and together are proposed to form a ‘map’ of the environment^16–18^. This code is proposed to be different for each distinct spatial context encountered, as evidenced by ‘remapping’: where the neurons that are active in a particular location in one context are not in the equivalent location in another^17–19,32^. Together this provides a representation of cues and events that is distinct dependent on the context in which they were experienced, allowing for context-specific behavior^33^. We therefore hypothesized that vCA1 neuronal activity might differentiate the abstract contexts required for performance of the 2-arm bandit task in a similar way to the well documented changes across spatial contexts.

We recorded the activity of neurons in the ventral two thirds of the CA1 area (vCA1), as the afferent and efferent connections of this part of hippocampus are heavily associated with reward and outcome representations^34^, and it is implicated in many goal directed behaviors^35^. We used microendoscopic imaging of single neuron Ca^2+^ activity in expert mice while they performed the 2-armed bandit task. We imaged a total population of 592 vCA1 pyramidal neurons across 6 sessions from 3 mice while they performed the task (**Figure 4A,B**).

**Figure 4.**
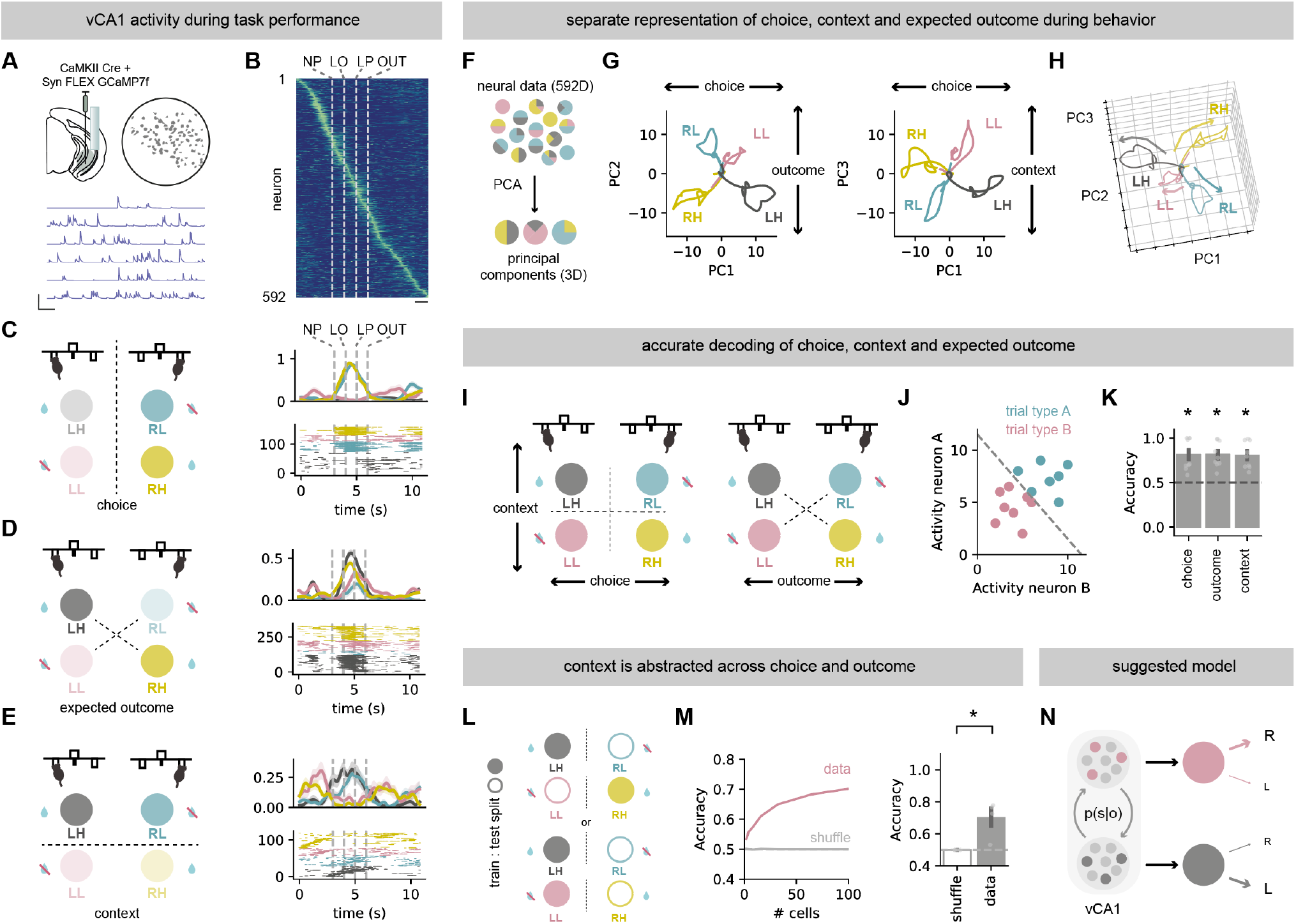
vCA1 maintains a generalized representation of abstract context during task performance. **A**. Top, schematic of viral injections and GRIN lens placement for Miniscope recordings of vCA1 pyramidal neurons in mice during 2-armed bandit task (left) and example field of view from one recording session (right). Bottom, example activity traces of individual neurons. Scale bar = 2Z, 20 seconds. **B**. Average activity of all recorded neurons. Scale bar = 1 s. **C**. Left, schematic of trials split according to the choice side. Right top, average spike probability of example vCA1 neuron across each trial type. Right, bottom, inferred spikes of the same example neuron. Note firing is consistently only on right choices, irrespective of anticipated outcome. **D**. Same as **C**, but for a neuron selective to high reward probability choices across both lever identities. **E**. Same as **C**, but for a neuron selective to either choice in a specific context – both left high probability (LH) and right low probability (RL) choices. **F**. Schematic of principal component analysis (PCA) of recorded neural activity. **G**. Top 3 principal components demonstrating separable representations of choice (left or right lever press, PC1), expected outcome (high or low reward probability choice, PC2), or latent context (current block of choice and reward contingencies, PC3). **H**. 3D representation of three PCs. **I**. Schematic of different task variable splits used in decoding analysis: choice (RH and RL vs LH and LL), expected outcome (RH and LH vs RL and LL), and latent context (RH and LL vs RL and LH). **J**. Schematic of SVM analysis. **K**. Decoding accuracy of animals’ choice, expected outcome or latent context from activity of simultaneously recorded vCA1 neurons. **L**. Schematic of a train:test split for decoding of generalized state representation – training performed on one of two conditions in each state (e.g. LH and RH) and then the decoding accuracy tested on the remaining two conditions from the corresponding state (e.g. RL and LL). **M**. Accuracy of decoding generalized latent context using either neural activity from a pseudo population of between 1 to 100 randomly selected neurons from all sessions from all mice (left) or in individual sessions with over 100 simultaneously recorded neurons (right). **N**. Schematic of a proposed latent context representation of a 2-armed bandit task by vCA1 pyramidal neurons. Points indicate individual sessions where unique neurons were recorded. Error bars represent s.e.m. across sessions in 3 mice (n = 9).

In the 2-armed bandit task, in contrast to exploration of dispersed spatial locations, mouse behavior is concentrated around one wall of a small operant chamber. As a result, instead of navigating through a maze, mice progress through different stages of each trial – from nose poke, to lever press to outcome while remaining in approximately the same spatial location. These trials can then be split into 4 types – either right or left choice, and whether that lever is currently associated with a high or low probability of reward delivery (abbreviated here and throughout as right-high: RH (yellow), right-low: RL (cyan), left-high: LH (black), left-low: LL (magenta)). A full representation of the task therefore relies on differentiating three separate organizations of these 4 trial types: i) split by choice (right versus left), ii) split by expected outcome (high versus low probability irrespective of choice), and iii) split by context (right high and left low – ‘context A’, versus right low and left high ‘context B’ – a readout of the contingency of the task).

First we looked for activity of individual neurons that was tuned to specific features of the task. As has been previously reported we found multiple neurons that were active only at specific times across the trial, irrespective of choice or outcome (**Sup. Figure 4**). However, we also found a large proportion of neurons that were active only on specific trial types (**Sup. Figure 4**). For example, activity of distinct neurons was separated across choice (∼40% of recorded neurons, e.g. **Figure 4C**), across the expected outcome irrespective of choice (∼20%, e.g. **Figure 4D**), but also the contingency of the levers, irrespective or either choice or expected outcome (∼30%, e.g. **Figure 4E**) – a representation of the latent context. Thus, similar to ‘place cells’ that tile locations and cues, and differentiate spatial contexts; there are also vCA1 neurons that tile the different parameter spaces of the 2-arm bandit task, and differentiate the latent contexts utilized to solve the task.

We next asked how the population as a whole represented each of these variables. To do this we first took the average activity of each neuron for each of the 4 trial types, and projected this onto the low dimensional space that captures the greatest variance across the different trial types (**Figure 4F-H**). We found that the top three components (PCs) almost perfectly separated trials based on choice (PC1), expected outcome (PC2) or context (PC3). Across these PCs, activity during each trial type was highly separable – suggesting rich representations of each trial type and task contingency in hippocampal activity. Importantly this same organization was found in individual mice, as well as the overall population (**Sup. Figure 4**).

We hypothesized that this organization should result in robust readout of each task variable. To test this, we designed a series of linear decoders to ask to what extent choice, expected outcome and context could be decoded from neural activity from each behavioral session from each mouse, limited to periods before the outcome of each trial (**Figure 4I,J**). Using this analysis, consistent with the organization of population activity, we found that choice, expected outcome and context could all be reliably decoded from neuronal data (**Figure 4K**). Moreover, by repeating our decoding analysis using small 1 s epochs spread evenly across the trial period, we found that these variables could be decoded stably across each trial and even during the ITI (**Sup. Figure 4**). Together this suggests that consistent with the role of vCA1 to support the use of SI during the 2-armed bandit task, neurons in vCA1 form a detailed, stable representation of choice, the expected outcome associated with that choice, and importantly the latent context of the task in which that choice is being made.

### Representations of latent context in vCA1 abstract across choice and outcome

Our results so far suggest that there is a representation of latent context in vCA1. This representation of context could occur via two means. First, individual neurons may represent the interaction between choice and expected outcome. For example, a neuron may fire only on RH trials (a trial with a right choice only when the right choice is high probability). Alternatively, vCA1 may contain a more abstract representation of state – where neurons have generalized representations of the latent context, irrespective of the trial type that is currently being performed. Such representations would be similar to a stable, ‘state’ representation often proposed to be the basis of contextual associative learning^4–10^.

We found neurons that exemplified both of these scenarios, with individual neurons firing to only one of the 4 trial types (**Sup. Figure 4**), and other neurons that had similar firing patterns across the two distinct trial types within each latent context, but different firing patterns across latent contexts (**Figure 4E, Sup. Figure 4**). To more quantitively test the presence of this more general representation, we built a separate series of decoders that were trained only on one trial type in each latent context (e.g. RH vs LH, or RH vs RL, **Figure 4L**). These models were then tested for their ability to accurately decode the other option in the same context (e.g. in an abstracted representation, training on RH trials should be able to correctly predict LL trials in testing). Using this approach, we found that vCA1 activity could decode these abstractions well above chance, both in overall population activity, and on a session-by-session basis (**Figure 4M**). Together this suggests that vCA1 represents choice, expected outcome, and latent context information, but crucially contextual information is abstracted and generalized across choice and outcome. Overall, our data are consistent with a model where latent context is represented in vCA1. This stable representation is ideally placed to be used by downstream areas such as prefrontal cortex, orbitofrontal cortex and NAc to define optimal behavior in each context^1–3^ (**Figure 4N**).

## DISCUSSION

The ability to use subjective experience to guide contextual decision making is fundamental for everyday life, but the neural basis of this ability has remained elusive. In this study we show that the hippocampus -an area strongly associated with spatial contextual representations - supports decision-making utilizing non-spatial, latent contextual representations^36,37^ (**Figures 1-3**). We found that neural activity in vCA1 robustly and stably differentiated two latent contexts formed only from past probabilistic outcomes (**Figure 4**) in a manner similar to that utilized to differentiate contexts encountered during spatial navigation^19,38,39^. Based on the large literature suggesting a key role of the hippocampus in spatial contextual associations^17,18,20,32,38^, this suggests a core function for the hippocampus may be in supporting the differentiation of, and inference across, contexts.

Much recent investigation of the basis of hidden state inference has focused on the role of the frontal cortex (FC), including orbitofrontal cortex (OFC) and medial prefrontal cortex (mPFC)^40^. Neurons in both rodent and primate FC have been shown to have strong representations of latent contexts combined with the value associated with different cues and actions within these contexts^3,41,42^, and activity in these regions is required for signatures of SI to be present in NAc dopamine release^43^. It is often presumed that the FC inherits contextual representations from the hippocampus, and uses this information to plan and assign value to cues and actions in each context^1,2,40^. Indeed, it has been shown that inactivation of ventral hippocampus impairs the representations of latent context in OFC during a similar reversal learning task^44^, but the nature of this representation has never been investigated. vCA1 projects strongly to mPFC, and thus indirectly to OFC^34,44^, with specialized connectivity that provides an ideal basis for tight control of FC circuitry by vCA1^45^. In this study we show that inhibition of vCA1 impairs both the dopaminergic and behavioral use of latent contexts to perform hidden state inference (**Figure 3**), and reveal the neural representations that underlie this key role (**Figure 4**). Future work will investigate whether this influence on dopamine signaling and behavior is via vCA1 projections to FC, or via more direct connectivity such as the strong projections from vCA1 to NAc^46–51^.

Finally, the hippocampus is a key node in some of the most debilitating mental illnesses including generalized anxiety disorder, depression and schizophrenia^52–54^. However, the neuronal basis of the role of the hippocampus is still much debated^11,12,33,55,56^. In this study we define a key role for hippocampal circuitry in the use of hidden state inference during reversal learning, an ability that is associated with the incidence and severity of a number of mental illnesses^11–14,57^. Future work will investigate how hippocampal dysfunction associated with risk factors for these illnesses disrupt the use of this strategy to guide behavior.

## METHODS

### Animals

6-9 weeks old (adult) male C57BL/6 mice provided by Charles River were used for all experiments. Animals underwent stereotaxic surgery and returned to their home cage for at least 1 week to allow full recovery. Animals were housed in cages of 1 to 4 and kept in a controlled environment under a 12h light/dark cycle with *ad-libitum* access to food and water (unless stated otherwise). All experiments followed Home Office and University College London guidelines.

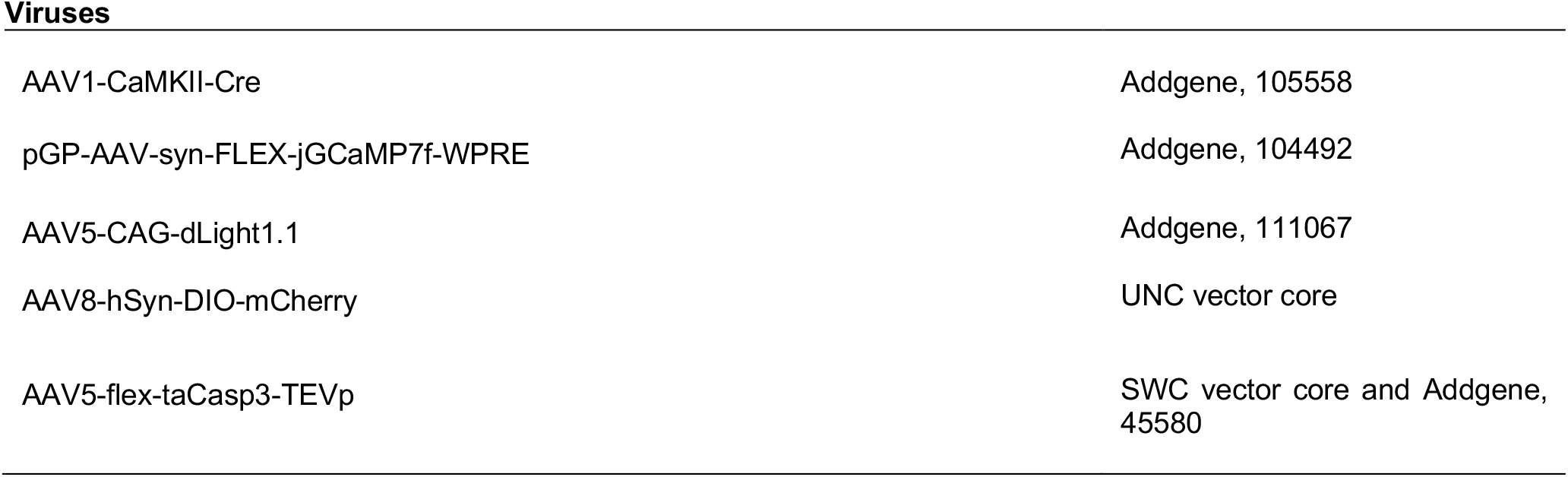

### Stereotaxic surgery

Stereotaxic surgeries were carried out according to previously described protocols^34,45–47^. For induction, mice were placed in a red perspex chamber (AN010ASR; VetTech) with 1.75 L/min flow of 4% vaporized isofluorane (in medical oxygen, 99.5% minimum purity). Following induction, fur on the scalp was shaved off using a small trimmer (ChroMini Pro; MOSER), and the animal was secured onto a stereotaxic head frame (Model 902 Dual Small Animal Stereotaxic Instrument; KOPF). Mice were placed on a homeothermic blanket control unit which was maintained between 35 and 37°C throughout the surgery (50-7001; Harvard Apparatus). During induction and throughout the surgery, the induction chamber and the stereotaxic frame were connected to an activated carbon scavenging filter (Cardiff Aldasorber; Shirley Aldred & Co) and an active scavenging unit (Model AN005; VetTech). For the duration of the surgery anaesthesia was maintained at the same flow rate and isofluorane concentration of 1-2%. Ophthalmic ointment (Viscotears® Liquid Gel) was applied to the eyes. The scalp was sterilized with HiBiSCRUB® and the skull was exposed with a single incision along the midline followed by application of a local anesthetic (0.025% Marcaine). After removing the connective tissue with sterile cotton buds, small holes were drilled in the skull at the coordinates of interest using a stainless steel bur (19008-07; Meisinger) attached to a miniature drill (Ideal Micro-Drill®; CellPoint Scientific). Injections were carried out with a Nanoject II (Drummond Scientific) using borosilicate glass pipettes back-filled with mineral oil and front-filled with ∼1 µL of the substance to be injected. 120 to 500 nL of virus was injected at a rate of 200 nL/min. Following infusion of the virus, the pipette was left in place for an additional 5 minutes before being slowly retracted. Injection coordinates were as follows (mm relative to bregma):

| Region | ML | RC | DV |
| --- | --- | --- | --- |
| Nucleus accumbens (NAc) | ± 0.9 | + 1.1 | – 4.6 |
| Ventral hippocampus (vH) | ± 3.2 | – 3.7 | – 4.5 |

After injection, the wound was sutured and sealed. Mice were given a subcutaneous injection of carprofen (0.5 mg/kg) and allowed to recover for a minimum of 30 minutes in a heated chamber before they were returned to their home cage. Animals received carprofen in their drinking water (0.05 mg/mL) for 48 hrs port-surgery.

For photometry experiments, mice were intracranially injected with 200 - 400 nL of *AAV5-CAG-dLight1*.*1* in NAc. For combined midbrain dopamine photometry and vH genetic lesion experiments, 200 – 400 nL of a 1:1 mix of *AAV1-CaMKII-Cre* and either *AAV8-hSyn-DIO-mCherry* or *AAV5-flex-taCasp3-TEVp* was injected into vH in the same hemisphere as NAc *dLight1*.*1* injection. Fiber optic cannula (200 µm core diameter, 0.39 NA, 5 mm long; Thorlabs) were implanted unilaterally above NAc following virus injection in the same surgery. To aid cement attachment, the skull was roughened, and two metal screws were inserted into the skull. Fiber implants were secured to the skull by applying two layers of adhesive dental cement (Superbond C&B). The skin was attached to the cured dental cement with Medbond skin glue (Animus).

For bilateral genetic lesion experiments, 200 – 400 nL of a 1:1 mix of *AAV1-CaMKII-Cre* and either *AAV8-hSyn-DIO-mCherry* or *AAV5-flex-taCasp3-TEVp* was bilaterally injected into 4 regions spanning the entirety of vCA1. The wound was sutured (6-0 Coated VICRYL polyglactin 910 suture; ETHICON) and sealed with Medbond skin glue (Animus).

For miniature microscope (UCLA Miniscope, Open Ephys) experiments, surgeries followed previous procedures^58^. Briefly, 1 – 2 mm diameter craniotomy was drilled at the vCA1 stereotaxic coordinates and the cortical tissue and corpus callosum fibers were aspirated using a blunt needle connected to a vacuum pump. Sterile saline (BAYER) was applied throughout aspiration to prevent desiccation of the tissue. 400 - 600 nL of a 1:1 mix of AAV1-CaMKII-Cre and pGP-AAV-syn-FLEX-jGCaMP7f-WPRE diluted in 2 parts of sterile saline solution (BAYER) was injected into vCA1. This dilution protocol was used to limit excessive GCaMP7f expression, which could lead to reduced Ca^2+^ variance in the signal, affect cellular processes and reduce cell health^58^. To increase the spread of the virus throughout the CA1/subiculum region of the vH, 3 injections of ∼165 nL each were delivered at –4.3, –4.5 and –4.7 DV coordinates. Relay gradient refractive index (GRIN) lens (0.6 mm diameter, ∼7 mm length, PN 130-000150; or 1 mm diameter, ∼4 mm length, PN 130-000143; Inscopix) were implanted either in the same surgery following injection of the viruses or 4 - 6 weeks after the initial surgery fixed to the custom-made base plate (Miniscopeparts) attached to the Miniscope for fluorescence guided implantation. The GRIN lens was inserted at an approximate rate of 0.5 mm/min to a depth between 3.5 – 4.3 mm and secured in place with super glue and further fixed with adhesive dental cement (Superbond C&B). To aid cement attachment, prior to the lens implantation the skull was roughened, and two metal screws were inserted into the skull. The Miniscope base plate was attached with adhesive dental cement (Superbond C&B) above the lens implanted directly to the skull. The base plate was locked to a Miniscope to find the optimal focus in the field of view prior to cementing. A protective cap was attached on top of the base plate to prevent debris build-up.

### Anatomy

#### Histology

Mice were anaesthetized with 0.5 - 1 mL of a mixture of ketamine (100 mg/kg; KetaVet) and xylazine (10 mg/kg; Zoetis) in sterile saline (BAYER). Following confirmation of deep anesthesia, animals were transcardially perfused with ice-cold 4% paraformaldehyde, the brains were dissected and fixed in 4% paraformaldehyde overnight at 4 °C. Brain samples were transferred to phosphate buffered saline (PBS, pH 7.2) after overnight fixation. Coronal brain slices were prepared at 70 µm using a vibratome (Campden Instruments). Slices were then mounted on gelatin-coated Superfrost glass slides with ProLong Gold, ProLong Glass Antifade Mountant with NucBlue (Molecular Probes), or Mowiol mounting medium. Fluorescent images were obtained with a 10x objective using a Zeiss slide scanner Axio Scan.Z1 using a 10x air immersion lens and standard filter sets for excitation/ emission at 365-445/50 nm, 470/40-525/50 nm, 545/25-605/70 nm and 640/30690/50 nm.

#### Immunohistochemistry

Brain slices (70 µm thick) were prepared as above and stained using standard procedures. First, slices were incubated in blocking solution (3% bovine serum albumin, 0.5% triton in PBS) for 1.5 - 3 hours at room temperature (22 – 24 °C) with constant agitation. When using the primary antibody raised in mouse, to eliminate non-specific binding, sections were first incubated overnight at 4 °C with anti-mouse F(ab)’2 Fragment in blocking solution and then washed 3 times for 20 - 40 minutes each in PBS. All slices were incubated overnight at 4 °C in blocking solution containing either 1:1000 anti-GFP (ab13970, Abcam) to reveal dLight1.1-expressing cells in NAc, or 1:500 anti-NeuN (Sigma-Aldrich, ZMS377) and 1:500 anti-GFP (Dako, GA524) to estimate caspase-induced cell loss in vH. Slices were then washed 3 times for 20 - 40 minutes each wash in PBS before incubation with secondary antibody(s) in blocking solution for 2 - 4 hours at room temperature (Alexa 647-conjugated donkey anti-chicken, AP194SA6, Millipore – to label GFP; Alexa 488- or Alexa 647-conjugated donkey anti-rabbit, A21206 / A31573, Invitrogen – to label GFAP; or Alexa 488- or Alexa 555-conjugated donkey anti-mouse, A21202/ A31570, Invitrogen – to label NeuN). Slides were mounted after a further 3 washes in PBS as above.

### Probabilistic reversal learning task

#### Behavioral setup

We trained animals on a probabilistic reversal learning task^23^. Following a minimum of 7 days of recovery after surgery, mice were water-restricted to approximately 85% of their ad-libitum weights. After at least a week of water-restriction and habituation to manual handling by the experimenter, behavioral training for the probabilistic reversal learning task began. All behavioral experiments were performed in 21.59 x 18.08 x 12.7 cm modular operant chambers (MED Associates, ENV-307W). Each chamber was equipped with a stainless-steel grid floor, two stainless steel walls (front and back), and a transparent polycarbonate side-wall, ceiling, and door. The nose port and the stainless steel reward delivery spout were located in the middle of the front wall. Two retractable levers were located either side of the nose port on the front wall (spout placed above the nose port). The behavioral box was also equipped with a house light placed outside of the chamber. Auditory stimuli were presented to animals via a speaker located on the back wall. Experimental events were controlled and recorded using custom scripts in MED-PC IV software.

All training and recording sessions were 1 hour long. All events within a trial (nose poke, levers out, lever press, and reward delivery) were separated by a temporal delay drawn from a random distribution from 0.1 to 1 s in 0.1 s intervals. In all stages of the task, a successful lever press always triggered retraction of the levers. Rewarded trials were signaled by 0.5 s of 5 kHz pure-tone auditory stimulus (CS+) and the delivery of 6 µL of reward (10% sucrose in water). Reward omission was signaled by 0.5 s of white-noise (CS-). The beginning of each trial was signaled by the illumination of a nose port. All trials were separated by a constant 3 s intertrial interval which began at the end of CS.

#### Training stages

Prior to data collection, mice went through several stages of training. In the first stage, mice were presented with both levers and had to press either of them to obtain a drop of sucrose solution until they had made over 100 lever presses in a single session. The next stage required mice to learn to nose poke into the central port to initiate presentation of a lever (alternating across trials) that they had to press for reward. Mice were then trained to remain in the nose port for 200 ms: starting with 0 ms, the nose poke duration required for the levers to come out incrementally increased by 10 ms every 10 trials until it reached 200 ms. Following completion of over 100 trials in a single session, mice then progressed to the deterministic reversal learning task. In this training stage, both levers were presented simultaneously but in a given block of trials pressing only one of the two levers would result in a reward. The identity of the rewarded lever reversed after 10 to 32 rewarded trials. After 3 successive sessions of receiving over 100 total rewards and choosing the rewarded lever over 60% of the trials, the mice progressed to the full probabilistic reversal learning task. In the full version of the task, in a given block of trials, one lever was associated with 70% reward probability following a press (high-probability lever) while the opposite lever was rewarded with 10% probability (low-probability lever). The identity of the rewarded lever reversed after 10 to 32 high-probability lever choices. Animals were trained on the full version of the task until they reached the ‘expert’ level with the consistent performance of over 60% high-probability lever choices for 3 consecutive sessions. While training for either the deterministic or probabilistic stage of the task, mice were also habituated to having optic fibers attached to the implanted ferrules or carrying a dummy Miniscope attached to the implanted baseplate until they met the performance criteria of the corresponding stage. Miniscope or photometry recording experiments commenced after mice were fully habituated and met the performance criteria for the final stage of the task.

### Behavioral analysis

To estimate the number of trials taken by mice to switch their choices to a different lever after reward contingencies reversal, we fit the exponential curve to animals’ reversal behavior (proportion of high probability lever presses following the reversal). The fit then allowed us to directly estimate the number of trials taken before animals started choosing the new high probability lever 50% of total lever presses after reversal.

To quantify the influence of the past choice and reward history on animal’s choice on the current trial, we used a logistic regression model^21,23,25,59^. In this model, the target variable was represented by the probability of the current choice being the right lever press (C(i), 1 if right choice, 0 if left choice). Predictor variable consisted of 3 types of trial history regressors: R(i – j) is the rewarded choice history on trial i – j (1 if rewarded right choice, –1 if rewarded left choice, 0 otherwise), N(i – j) is the unrewarded choice history (1 if unrewarded right choice, –1 if unrewarded left choice, 0 otherwise), C(i – j) is the outcome-independent choice history on trial i – j (1 if right choice, –1 if left choice, 0 otherwise). The encoding model is:

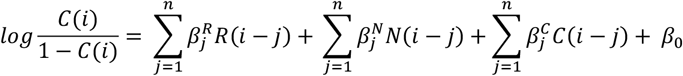

Where 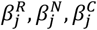 are the regression weights of each history predictor, and β_0_ is the history-independent constant bias term. While this regressor set is not strictly orthogonal, due to the inclusion of a separate choice predictor, it provides a biologically informed estimate of the contribution of choice and outcome on upcoming choice^25^. To model the animal’s choice given its trial history, the regression coefficients were fit using LogisticRegression function of *scikit-learn* Python library. For this model we used elastic net regularization, method that combines L1 and L2 regularization penalties. First, we performed grid search over *C* (inverse of regularisation strength) and l1-ratio (contribution of L1 and L2 penalties in the elastic net regularisation) hyperparameter space to find the optimal combination of *C* and l1-ratio that explained the most variance when verified with 5-fold cross-validation. The overall total explained variance of the final model *R*^2^ was calculated as an average from 5 cross-validated fits of the model with the best estimated *C* (0.25 ± 0.04) and l1-ratio (0.34 ± 0.04) hyperparameters. *C* and l1-ratio that provided the best average *R*^2^ score were then used to refit the full data set to obtain estimated weights *c*. The logistic regression coefficients were fit separately for each session in each animal. Estimated coefficients represented the extent the different past trial choices and outcomes predicted animals’ current choices. Model β coefficients were then used to estimate how much different past trial history predictors influenced animals’ decisions on the current trial.

### Behavioral Modelling

We investigated behavioral strategies mice might use when solving the probabilistic reversal learning task by fitting a range of different computational models to their choices. We considered a number of ‘simple’ models such as random choice, win-stay-lose-shift (WSLS) and choice repetition^60^ as well as more complex value updating and state inference strategies.

### Simple behavioral models

Random choice model assumes that mice do not engage with the task and press levers at random with a bias (*b*) for one option over the other. The probabilities of choices *a* and *a*′ on trial *t* is:

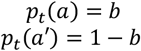

Noisy WSLS model repeats rewarded actions and switches away from unrewarded actions with probability 1 – *ε* and chooses randomly with probability *ε* The probability of choosing option *a* is:

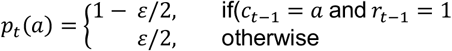

where *c*_t_ is the choice at trial *t*, and *r*_t−1_ the reward at trial *t*.

Choice kernel model tries to capture the tendency for mice to repeat their previous actions. Specifically, the agent computes a ‘choice kernel’, 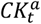, for each action, which keeps track of how frequently that option was chosen in the past.

The choice kernel updates according to the rule below:

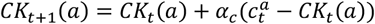

Where 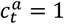 if lever *a* is chosen on trial *t*, otherwise 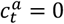, and *α*_c_ is the choice kernel learning rate. In the choice kernel model each option is chosen according to a softmax function:

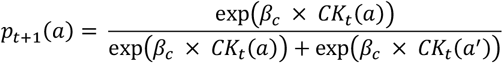

where β_*c*._ is the inverse temperature associated with the choice kernel.

### Value updating (Q) models

Value updating models are reinforcement learning (RL) models that utilize Q-learning updating rule. In such models on every trial *t* the expected value *Q*_*t*_(*a*) of action *a* is updated by the reward prediction error (RPE), difference between the choice outcome *r*_*t*_ and previous expected value *Q*_*t* −1_ (*a*), scaled by the learning rate *α*:

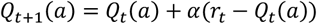

The choice probabilities were estimated based on the action values according to a softmax function:

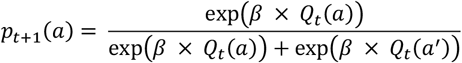

where β is the inverse temperature.

Other models from the reinforcement learning family were augmented alterations of the basic model above. Introducing bias captures animals’ preference towards one of the levers in the task.

Bias parameter *b* (−0.5 < *b* < 0.5) changes the expected value of one of the actions reducing or increasing the probability of choosing that action:

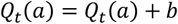

RL strategies may also be affected by animals’ tendency to repeat previously selected actions. To incorporate this into the model, we have added the choice kernel (described above) into the softmax decision rule:

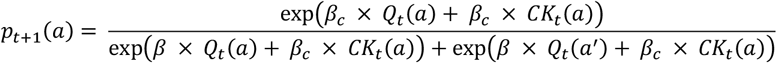

Reward and punishment (R/P model) sensitivity augmentation utilizes the same value updating rule while using different learning rates following rewarded (*α*_*r*_) and unrewarded (*α*_*ur*_) outcomes ^25,61^:

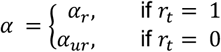

Models with reward sensitivity include the parameter *ρ* that allows to modify the effective size of reinforcement^12^:

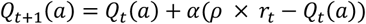

In double updating models, expected values for both actions are updated on every trial: values of the unchosen actions *a*’ are updated according to the counterfactual outcome (1 – *r*_*t*_) from the chosen action (*a*)^5^:

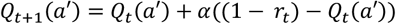

We tested three versions of the counterfactual updating models. In the ‘same *α*’ model, the learning rate *α* used for updating the action value of the unchosen option was the same as the learning rate used for updating the chosen option. Alternatively, the ‘different *α*’ model utilised different learning rate parameter(s):

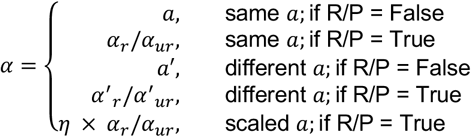

In value updating models with forgetting, expected value of the nonchosen action *a*’ was either directly reset over one trial to the average value *Q*_*t*_ across both actions (Forget reset) or gradually updated towards the average *Q*_*t*_ according to the forgetting factor *δ* (Forget gradual):

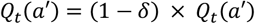

*Dynamic value updating models* are based on the basic RL strategy that utilizes the Pearce-Hall rule^62,63^. These models contain an associability parameter that modulates the learning rate as a function of the absolute magnitude of past RPEs. The *κ* parameter modulates the action value updating and is equivalent to the learning rate parameter in the basic RL models. The *γ* parameter controls the temporal dynamics of associability over time:

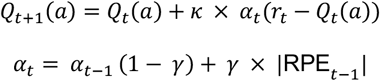

Like other value updating models described above, dynamic value updating models could also be modified to include bias, perseverance, and R/P as described above. R/P is enabled by having different *κ* parameters for rewarded and unrewarded trial outcomes:

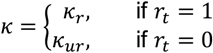

***State inference (SI) models*** assume that on each trial, mice chose their actions based on their belief *b*(*s*) about the underlying state of the task *s* given the observed outcomes following their past choices *o*_*t*_ *=* {*a*_*t*_, *r*_*t*_}.

The belief variable takes on a role similar to the Q value in the standard RL models above, becoming a function of the past observations and the parameters^12^:

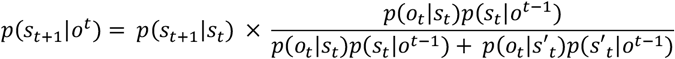

The most rigid instance of the state inference models utilizes the knowledge of the reward probabilities associated with each choice (Reward matrix, *Rm*) and the probability of block reversal on each trial^22^:

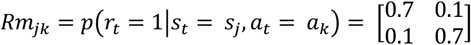

which is then used in calculating the probability of observations given state:

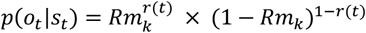

The 5% block reversal probability on each trial can be written in terms of the block state transition probabilities as (referred to as ‘fixed *γ*′ SI model):

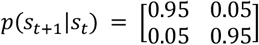

Finally, the current belief b = *p*(*s*_*t*_|*o*^*t*−1^) about the state of the task based on past observations is mapped into action probabilities via a softmax function as in the RL models:

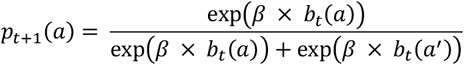

Regardless of the number of the states in this task, this model assumes that subjects only care about which of the two actions is best - that there are only two states corresponding to each action being the better one. Alternatively, we also tested models in which the action probability is estimated using the reward expectation in a given state based on the reward matrix *Rm* and current belief estimate *b*_*t*_:

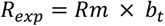

Next, we incrementally modified this rigid state inference model to make it more flexible. First, we have added a parameter *γ* that corresponds to a probability of staying in a state:

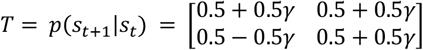

Next, we have added flexibility to the estimate of probability of an observation given the state. Instead of using experimenter defined reward probability matrix *Rm*, the outcome compatibility with the state was estimated. In this version of the state inference model, a reward following the choice *a* tells with probability *c* that it was a correct choice and that the current state is the state corresponding to action *a* being a high reward probability choice. Conversely, for the non-selected option *a*′ reward following choice *a* provides negative evidence for the choice *a*′ being a high reward probability choice with probability *c*.

Finally, reward omission provides following choice *a* provides evidence for choice *a*′ and against choice *a* being a better option. Similar to the R/P augment in the RL models, in the state inference model with reward and punishment sensitivity, different probability parameter *d* may be used following the reward omission.

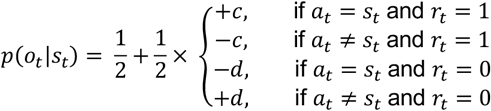

Because in the most fits to animals’ behavior the reward omission update parameter *d* was estimated to be very close to 0, we have also tested the model that only uses positive choice outcomes in updating the state prediction^64^, fixing parameter *d* at 0. Thus, following reward omission *p*(*o*_*t*_|*s*_*t*_) is 0.5 for both actions. We also tested versions of state inference models with the parameter β fixed at 10 ^12^.

### Model fitting

To estimate the values of the parameters that best describe the behavioral data, we used likelihood maximization approach to model fitting^60^. For this, for each behavioral session we estimated the probability of individual choices based on a given model (*m*), parameters of the model (Θ_*m*_) and choice and outcome history in that session. We then summed the logs of choice probabilities that corresponded to animals’ choices on every given trial. The python function *scipy*.*optimize*.*minimize* was used to find the set of parameter values that minimized the negative log of the likelihood of the data (*LL*, equivalent to maximising the likelihood) given the model parameters *p*(*d*_*1:T*_|Θ_*m*_, *m*):

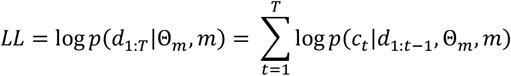

To avoid finding the local minima in the minimization procedure, we repeated model fitting procedure 50 times using randomly selected initial values from defined bounds for each parameter, and recorded the best fitting log likelihood for each run. The best fitting parameters were selected from the run with the highest log-likelihood value.

To determine which model provided the best fit to the data we compared different model fits using Bayesian Information Criterion (BIC). BIC has an explicit penalty for the number of free parameters (*k*_*m*_) in the model *m* and thus controls for overfitting:

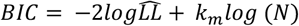

Where 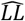 is the log-likelihood value at the best fitting parameter settings, and *N* is the number of trials. To compare the model fits for each animal, we then computed the differences between the BIC scores of the each model fit to individual behavioral sessions with the BIC score of the best fitting model to the same session (ΔBIC). The relative likelihood values of the model *m* compared to the best fitting model to the same session were calculated as follows:

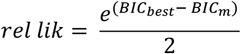

According to the BIC analysis, all behavioral sessions from all mice were best fit by SI models. Specifically, the best fitting SI models had reward and punishment sensitivity (R/P), used only rewarded trials for updates (*d =* 0), utilized the transition matrix with the probability of staying in a state *γ*, had parameter β fixed at 10; while some also had choice bias. Among the Q model group, the best fitting models either had only choice bias, or had both R/P and choice bias. Therefore, for our comparison of the SI and Q models in the main figures, we focused on the versions of best-fitting Q and SI models that included the same augmentations– R/P and choice bias. To assess changes in behavior in caspase lesion/sham animals (**Figure 3C-E**), mouse behavior was fit as above from sessions obtained at baseline before lesion. The effects of the lesion or sham were then assessed using these model parameters to investigate how model predictions of trial-by-trial behavior were alter by the manipulation.

### Model simulations

To simulate the probabilistic reversal learning behavior, we ran the models with the parameter sets obtained from the model fits to individual mouse sessions. Each set of parameters was used for 3 simulation runs, and simulation run comprised of 300 trials.

### Model verification

To check how reliably we can conclude that the best model from the fitting procedure was more likely to have generated the data compared to other models that were tested we performed model recovery^60^. For this, we used simulated data from all models and fit that data with all models. From this we quantified the proportion of the simulated data generated by one model that was best fit other models *p*(fit model|simulated model), summarized in a confusion matrix (**Sup. Figure 1**). If 100 % of the simulations were best fit by the same models that produced the simulated data, the confusion matrix would be the identity matrix. We also computed the inversion matrix that quantified the probability the model generated the data given that it provided the best fit *p*(simulated model|fit model), (**Sup. Figure 1**). From the inversion matrix we can estimate the confidence with which we can draw conclusions about the behavioral strategies based on the best fitting models (how likely the same model is to have generated the data).

### Photometry

#### Recording setup

To measure dopamine release, we recorded dLight1.1 fluorescence using a custom-built fiber photometry acquisition as described previously^45,47^. Briefly, to record dLight-dependent fluorescence we used blue 470 nm LED, while to control for dopamine-independent fluctuations in recorded fluorescence (e.g. due to movement) we used violet 405 nm LED. LEDs were controlled via a custom script written in LabView (National Instruments). To enable synchronization with the behavioral task, the recording was initiated by a TTL pulse from the MED-PC program at the start of the behavioral session. To ensure the separation of the blue and the violet channels, the light amplitudes were modulated sinusoidally with two different frequencies (500 Hz and 210 Hz, respectively). For excitation, light from both LEDs passed through corresponding excitation filters (470 nm and 405 nm) before being combined into a single beam by a dichroic mirror. The excitation light was then passed through a beam splitter to allow for simultaneous recordings in two animals. The excitation beams were then reflected off a dichroic mirror, collimated and launched into a fiber patch cord (200 µm core and 0.22 NA). The patch cord was connected to the ferrule of the implanted optical cannula on the animal’s head via an interconnect. The emission signal was passed through the same patch cord and collimator, and filtered through an emission filter (transmission above 505 nm). It then passed through a dichroic mirror and focused onto a femtowatt photoreceiver (Newport) sampling at 10 kHz. Each of the two modulated signals generated by the two LEDs was recovered using standard demodulation techniques implemented by a custom Labview script. dLight and control autofluorescence signals were then downsampled to 500 Hz before being exported for further analysis.

#### Photometry data processing

Photometry data were analyzed with custom-written Python scripts. First, to reduce the noise, a lowpass filter was used on both dLight and control signal. To correct for photobleaching, a 4th order polynomial fit was subtracted from each trace. The fluorescent signal obtained after stimulation with control 405 nm LED was used to correct for dopamine-independent changes in fluorescence such as due to movement. Movement artifacts were estimated by a least-squares linear fit of the control signal from 405 LED excitation to the dLight fluorescence. The estimated movement signal was then subtracted from the dLight signal to obtain the movement-corrected signal corresponding to dopamine release. Signals were then z-score normalized. For photometry experiments with chronic taCasp3 hippocampal inactivations, we excluded data from mice where dLight signals were not observable (2 mice from mCherry control group), or where mice had misplacement of caspase injections inferred from the immunohistochemistry labelling (2 mice from taCasp3 group).

#### Photometry data analysis

Dopamine signals were analyzed in two complementary ways. First, z-scored signals were aligned to CS and baselined to the mean signal from 1 s preceding the event. The event summary was obtained by calculating mean of the baselined signal in the first 4 seconds of the event. Selection of a wide time window for the event summary enabled us to capture most of the event-associated signal in an unbiased way irrespective of the temporal variability across animals. These events were then sorted according to past choice and outcome history (same or opposite choice, rewarded or non rewarded outcome), and compared to RPE calculated from simulations of agents utilizing Q or SI strategies. For Q estimates, RPE on a particular trial was the outcome minus the estimated value for that choice. For SI estimates, RPE was the outcome minus the reward estimated from the reward probability matrix. We compared model estimates qualitatively across pairs of choices and outcomes (fig, supp fig), but also investigated this more quantitively using a regression approach to predict dLight signal across each of the 8 trial types using either SI and Q RPE as predictors. Second, we used 2-fold cross validated ridge regression to express dLight fluorescence as a sum of responses related to outcome, past outcome, choice and past choice, as well as estimates of Q-RPE and SI-RPE from our model fits. For this analysis, photometry traces were aligned across trials by linearly time-warping the signal at the intervals between different fixed trial events and resampling the signal at a fixed rate. We only included data within 6 trials of a switch in contingency, due to the increased number of incongruent trials allow better discrimination of the two RPE predictors. Behavioral predictors (outcome, past outcome, choice and past choice) were binary variables centered at 0 (i.e. outcomes were coded as 0.5 for rewarded and -0.5 for non-rewarded), while latent variables (SI- and Q-RPE) were continuous estimates from model fits. For each time point we calculated the coefficient of partial determination (CPD) for each predictor, i.e., what percentage of the variance of the dLight activity at that time-point was explained by the full regression analysis that was not explained by the regression analysis if that predictor was removed. To compliment this we also performed single predictor regressions where we calculated the variance that could be explained by only one predictor. For both of these metrics we compared the contribution of SI- and Q-RPE to model fits, and the influence of vCA1 lesions.

### Miniscope

#### Recording setup

Calcium imaging was acquired using Miniscope V4 – a head-mounted microscope (OpenEphys) controlled via Miniscope-DAQ-QT-Software. A blue LED was used for excitation (∼470 nm spectral peak) with power adjusted to approximately match the mean brightness of the image across animals. Fluorescence was passed through an emission filter (bandpass filter, 525/50 nm) and collected by a CMOS imaging sensor. Before the start of the recording, the Miniscope was attached to the base plate and its focal plane was adjusted. Afterwards, the mouse with the Miniscope attached was placed in a behavioral chamber (MED Associates, ENV-307W) for 3–5 minutes before the recording session started. Miniscope was connected to a laptop via a flexible coaxial cable and an off-board data acquisition (DAQ) board and the calcium imaging data was acquired at 30 Hz using Miniscope-DAQ-QT-Software (https://github.com/Aharoni-Lab/Miniscope-DAQ-QT-Software#miniscope-daq-qt-software).

#### Miniscope data processing

Minian software was used for all pre-processing stages and subsequent fluorescence signal extraction^65^. To improve the computational performance of the processing pipeline, the videos were first cropped to a rectangle containing the imaged cells, the video width and height was downsampled by a factor of 2, and the framerate was downsampled by a factor of 2. Following the correction of the background fluorescence and median filtering for sensor noise removal, the video was motion-corrected and seeds for estimation of cells’ spatial footprints were initialized. This set of seeds was then used for cell and signal detection using a constrained non-negative matrix factorization (CNMF) algorithm^66^. Following the refinement of the spatial footprints and denoising of the temporal traces of each cell, the CNMF algorithm produced background-subtracted calcium fluorescence values and deconvolved the calcium trace into estimated ‘spikes’ that corresponded to a scaled probability of neural activity. The results were then visually inspected and non-cell like shapes and traces were excluded from the output. Deconvolved calcium traces were subsequently aligned across trials by time-warping as described for photometry above.

#### Selectivity index analysis

Trial type selectivity of individual neurons was computed as:

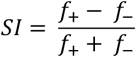

where *f*_+_ and *f*_−_ are the average activity of the neuron in the period from 1s before trial initiation up to the CS delivery on different trial types (right vs left, high or low reward probability, state A vs state B choices)^67^. To assess the statistical significance of selectivity indices (SIs) of individual neurons, we compared their SI values to those derived from 1000 shuffled datasets, where the labels of trial types were randomly reassigned.

#### Neuronal trajectory analysis

The neural population activity trajectories were obtained by projecting the average population activity for each trial type into the low dimensional space that captured most variance between trial types. Every trial in the task belonged to one of four conditions defined by the combination of animal’s choice and reward contingency associated with the chosen lever in a current block of trials: left-high (LH), right-high (RH), left-low (LL), and right-low (RL). First, to evaluate the component of activity that was not selective to different trial types, we calculated the average activity for each neuron across all trial types. We then subtracted the non-selective activity for each neuron from that neurons average activity for each individual trial type, baselined to isolate within-trial variation, and concatenated across trial types to generate a data matrix representing how activity for each neuron deviated from its cross-trial-type average in each trial type^68^. We performed PCA on this matrix to find the space that captured the most cross-trial-type variance and then projected the average population activity trajectory for each trial type into this space.

#### Population decoding analysis

The decoding analysis was used to predict different trial types based on mean spiking probability of simultaneously recorded neurons. For this, average neural responses associated with different task variables we estimated as a mean spiking probability of individual neurons from 1s before the trial initiation up until CS. Based on different combinations of animal’s choices and associated reward probabilities outlined above, each trial could be classified based on animal’s choice (lever identity – LH and LL vs RH and RL), expected outcome (reward probability associated with the chosen lever – LH and RH vs LL and RL) or task state (a set of reward contingencies associated with either choice in a given block of trials – LH and RL vs LL and RH). To balance the number of different trial types, for each neuron each class trial pool was randomly sampled 250 times with replacement. Unless states otherwise, the decoding analysis was performed on activity from simultaneously recorded neurons from a single behavioral session.

For decoding, we used a support vector machines (SVM) classifier with a linear kernel implemented through LinearSVC function from *scikit-learn* Python library. For cross-validation, data were randomly divided into two non-overlapping groups of trials, used for training and testing the classifiers (75/25% split). The decoder was trained to discriminate between population responses corresponding to two sets of trial variables representing either animal’s choice, expected outcome, or task state. The regularization hyperparameter C was optimized using GridSearchCV *scikit-learn* function with 5-fold cross-validation. The decoding performance was then tested using a held-out test set. This procedure was repeated at least 100 times for each classifier with random train/test subdivisions and the decoding accuracy was computed as the average result across repetitions. To assess the statistical significance of the decoding accuracy, we repeated the same procedure described above on a dataset with shuffled trial labels.

For cumulative decoding plots, we generated neural pseudo-populations from subsets of neurons sampled across multiple animals and/or recording FOVs. Each trial condition was sampled 250 times and activity of individual cells within that condition shuffled. The decoding analysis was performed as for single session models outlined above.

#### Generalization decoding analysis

To quantify the degree of generalized representation of the trial variables, we used cross-condition generalization performance^69^. In generalization decoding analysis training and testing sets were created by splitting trials according to their trial labels, so the decoder was trained to discriminate trial categories according to half of the labels and then the discrimination generalization was assessed on the data from different conditions not used in training. For example, to test generalized encoding of choice, the decoder was trained to discriminate RH and LH trials and its performance was tested on discrimination of RL and LL trials, respectively.

### Statistical analysis

All statistics were calculated using the Python packages *scipy, pingouin* and *statsmodels*, and *lme4* R package implemented in Python through rpy2. Summary data are reported as mean ± s.e.m. (standard error of the mean). Unless otherwise stated, statistical tests were performed comparing data from individual behavioral sessions, including mouse identity as a random effect to maintain the dependence between sessions from individual mice. As a result, degrees of freedom are estimated using the Satterthwaite approximation. Normality of data distributions was determined by visual inspection of the data points. Test statistics are detailed in the supplementary statistics table. Threshold for statistical significance was defined as 0.05. No power analysis was run to determine sample size a priori. The sample sizes chosen are similar to those used in previous publications. Throughout the figures the * symbol represents *p* < 0.05.

## ACKNOWLEDGEMENTS

We thank members of the MacAskill laboratory for helpful comments on the manuscript. A.F.M. was supported by a Sir Henry Dale Fellowship jointly funded by the Wellcome Trust and the Royal Society (grant number 109360/Z/15/Z) and an MRC project grant (MR/W02005X/1). K.M. was supported by the Wellcome Trust 4-year PhD in Neuroscience at UCL (grant number 215165/Z/18/Z).

## AUTHOR CONTRIBUTIONS

Conceptualization, Methodology, Investigation, Formal Analysis, Writing – Original Draft, Writing – Review & Editing, K.M. and A.F.M.; Funding Acquisition, Supervision, A.F.M.

## INCLUSION AND DIVERSITY

We support inclusive, diverse, and equitable conduct of research.

## SUPPLEMENTARY FIGURES

**Sup. Figure 1.**
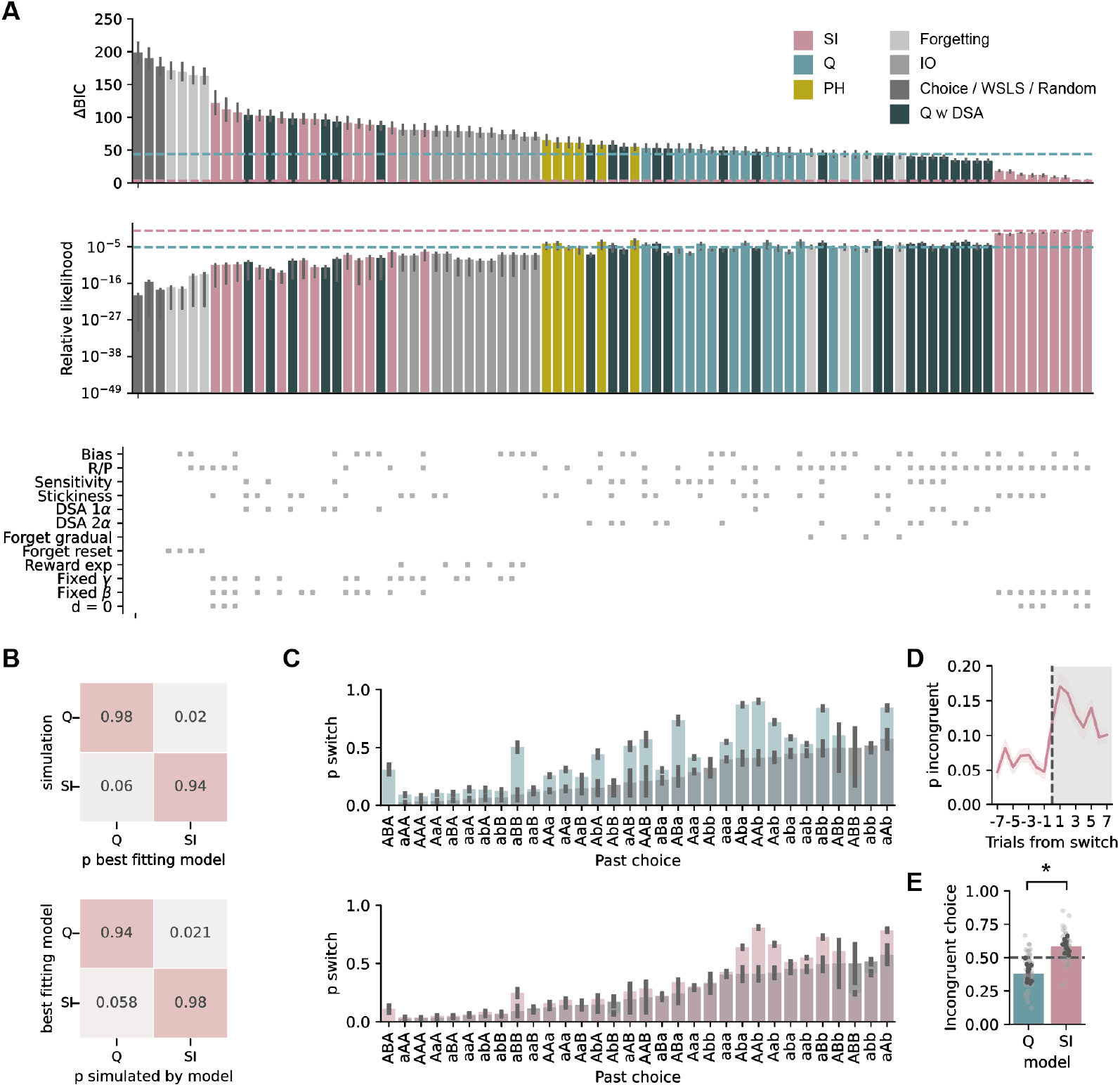
Model fits of mouse behavior. **A**. ΔBIC (top), and Relative likelihood (middle) of all model fits. Model types are color coded, and additions such as bias and stickiness are shown in raster plot (bottom). **B**. Confusion (top) and inversion (bottom) matrix showing Q and SI models are distinct and readily distinguishable. **C**. Mouse switching behavior (grey) split by past choice and outcome history (as shown in ^25^). Q (top, blue) and SI (bottom, pink) model predictions are overlayed, note SI prediction is most representative. **D**. Proportion of incongruent trials (trials where Q and SI models had different predictions) around a switch in contingency. **E**. Proportion of incongruent trials consistent with either Q or SI strategies.

**Sup. Figure 2.**
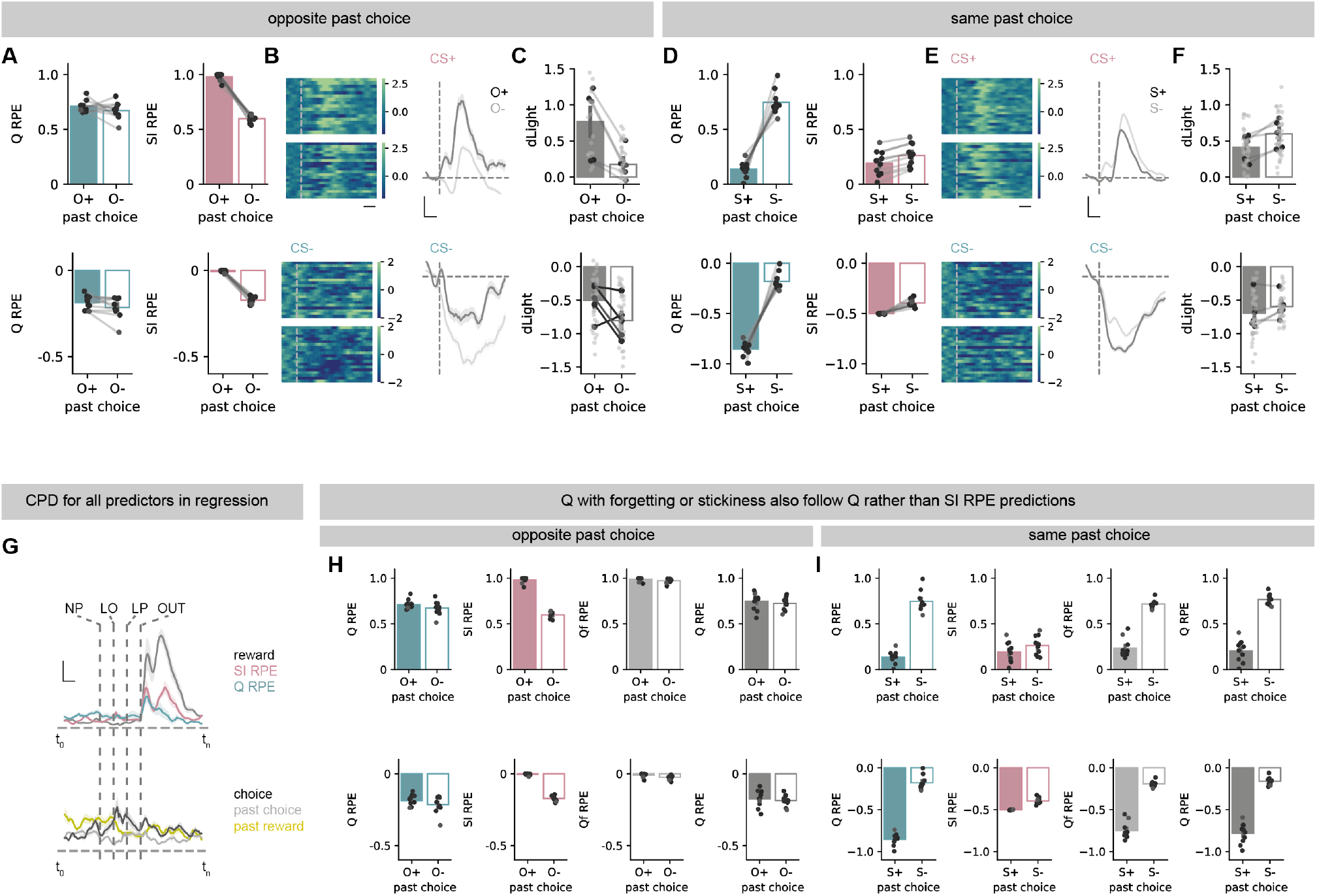
Dopamine dynamics across different choice and outcome histories. **A**. Prediction error estimates from Q (left) or SI (right) models for trials split according by outcome of past opposite lever press (see text): O+ (reward) or O-(reward omission). Top shows current CS+, bottom shows current CS-. Note only SI predictions change. **B**. Left, z-scored dLight signal aligned to CS for O+ (top) or O-(bottom) trials. Right, example z-scored dLight signal around CS for O+ and O-trials for one session. Scale bar = 1 s (left); 1s, 0.5 zF (right). **C**. Summary of CS dLight signal on O+ and O-trials. **D-F**. As (**A-C**), but split by the outcome of the same past choice. **G**. Coefficients of partial determination for top: outcome (grey), Q-RPE (blue), SI-RPE (pink), bottom: choice (black), past choice (grey) and past reward (yellow) at each timepoint. Scale bar = 1s, 2.5 %. **H-I**. Prediction error estimates as in **A** and **D** from Q (blue) or SI (pink) and Q with forgetting^27^ (light grey) and Q with stickiness (dark grey) models. Note that Q with forgetting and Q with stickiness have equivalent predictions as Q, and are distinct from SI. Dark points show individual mice, light points show individual sessions. Error bars represent s.e.m. across animals (n = 5).

**Sup. Figure 3.**
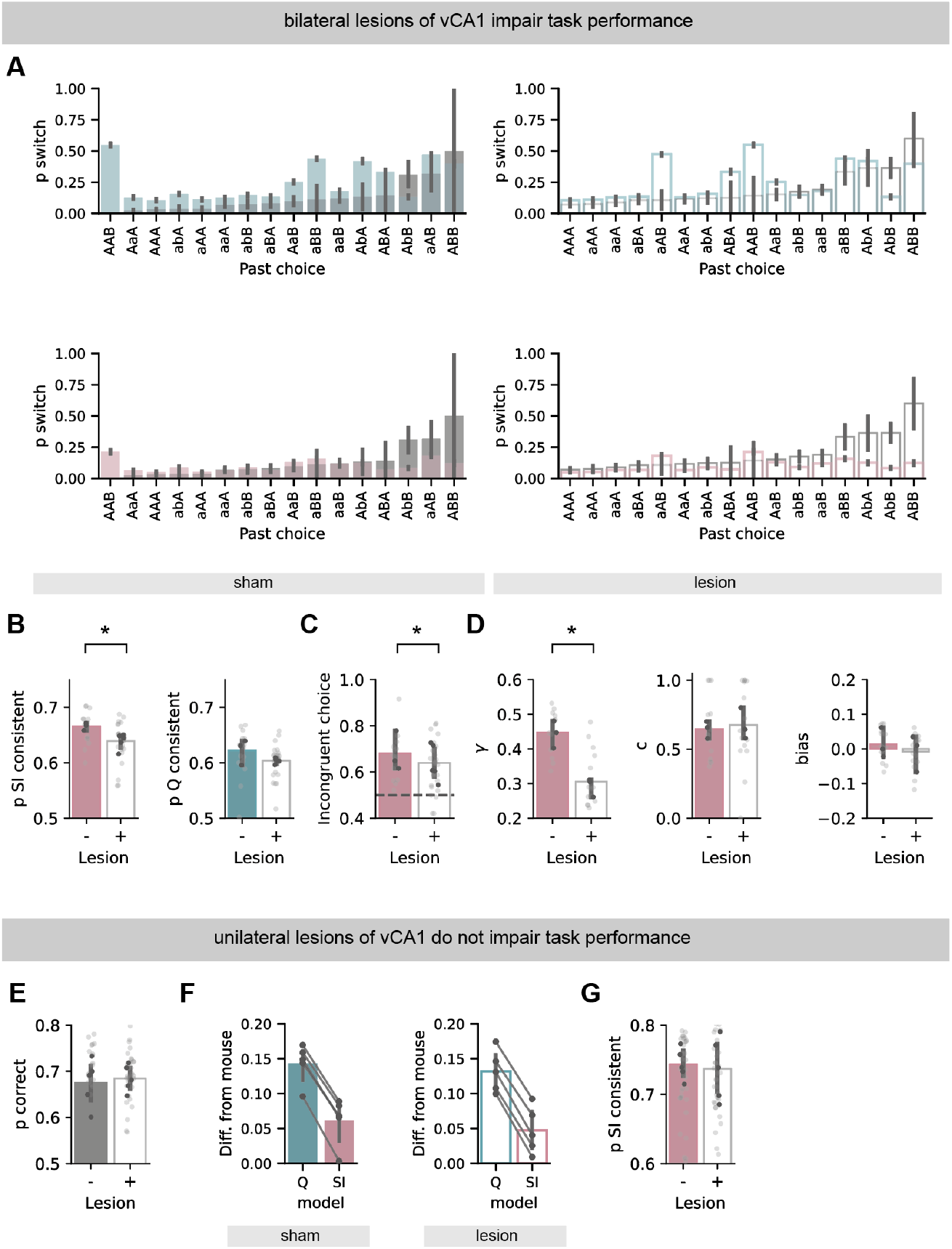
The influence of bilateral and unilateral vCA1 lesions on behavior. **A**. Mouse switching behavior split by past choice and positive outcome history. Q (top) and SI (bottom) model predictions are overlayed on sham (left) and lesioned (right) animals. Summarized in **Figure 3C,D. B**. Proportion of choices consistent with SI strategy (left) and Q strategy (right) in sham and lesioned mice around a switch in contingency. Note lesion has limited effect on Q strategy consistent choices. **C**. Proportion of incongruent choices consistent with SI strategy in sham and lesioned mice. **D**. SI model parameters for fits to sham and lesions sessions. **E-G**. Unilateral lesions do not affect behavior: **E**. Summary of high probability choices in sham and caspase lesioned mice. **F**. Difference between mouse switching behaviour at different trial histories, and that of simulations utilising either Q or SI strategies in sham and lesioned mice. **G**. Proportion of choices consistent with SI strategy in sham and lesioned mice. Dark points show individual mice, light points show individual sessions.

**Sup. Figure 4.**
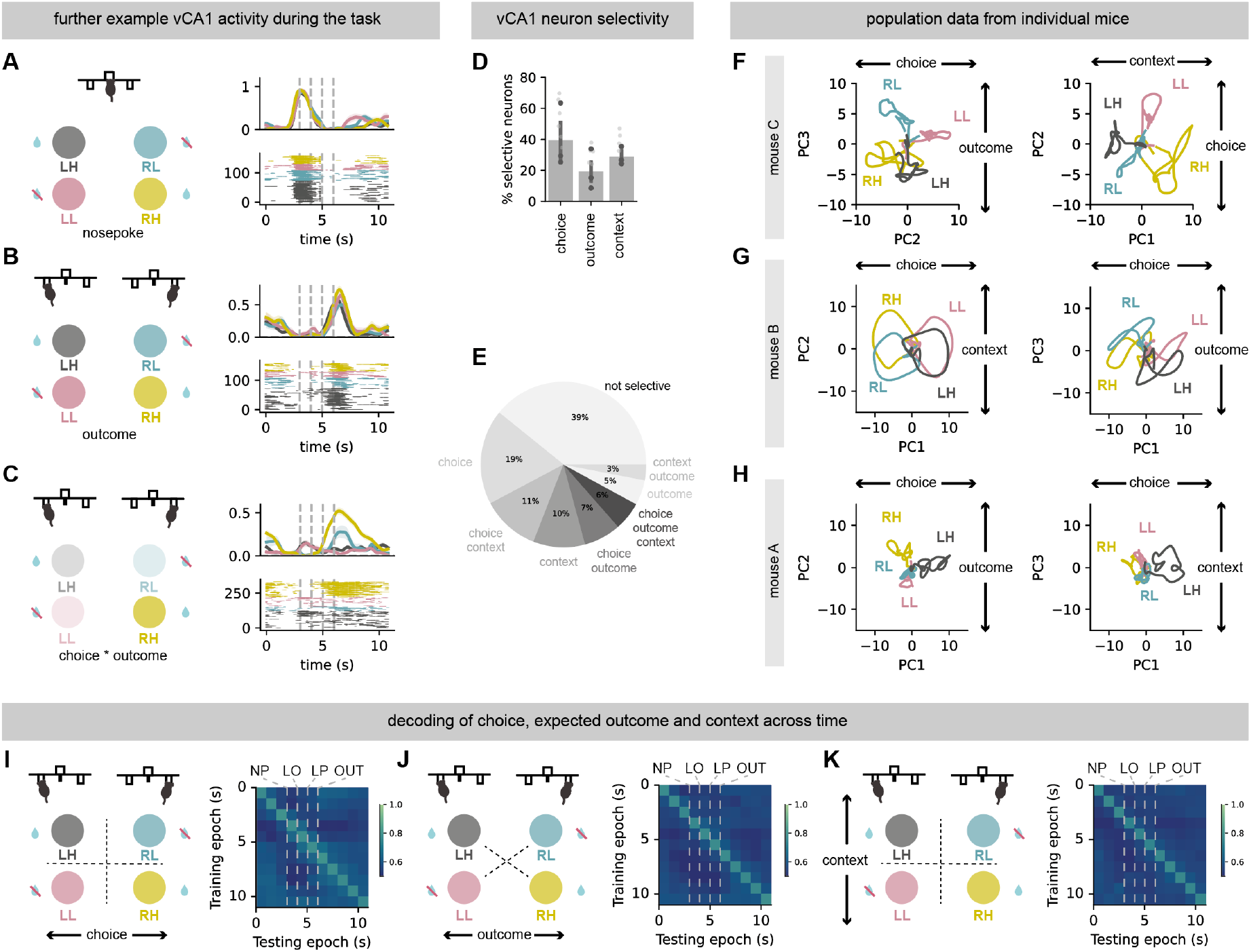
Analysis of vCA1 activity, trial selectivity and consistency across mice. **A**. Example vCA1 neuron with activity preferentially around trial initiation (nosepoke), irrespective of trial type. **B**. As in A, but neuron tuned to outcome, irrespective of trial type. **C**. Example neuron with activity preferentially on only RH trials. **D**. Summary of proportion of neurons selective for either choice, expected outcome or context. **E**. Proportion of neurons selective for different combinations of choice, expected outcome or context. **F-H**. Principal components calculated as in **Figure 4F**, but for individual mice. Note that each mouse has similar separation of choice, expected outcome and context across the first three PCs. **I-K**. Decoding analysis as in **Figure 4K**, for choice (**I**), expected outcome (**J**) and context (**K**), but for individual 1s epochs throughout the trial. Note that decoding of each variable is stable across the trial and ITI. Dark points show individual mice, light points show individual sessions.

## STATSTICS SUMMARY

STATISTICS SUMMARY
| Figure | Descriptors | n | Test used | Statistic | p-value |
| --- | --- | --- | --- | --- | --- |
| 1G | ΔBIC<br>Q vs SI fit | 10 mice | Paired t-test | $t_{(9)} = 8.76$ | 0.000011 |
| 1H | Relative likelihood of<br>Q vs SI fit | 10 mice | Paired t-test | $t_{(9)} = 12.7$ | $4.7 \times 10^{-9}$ |
| 1I | Difference between mouse<br>and prediction from Q vs SI | 10 mice | Paired t-test | $t_{(9)} = 9.73$ | 0.000005 |
| 1J | p choice consistent with<br>Q vs SI | 10 mice | Paired t-test | $t_{(9)} = 3.55$ | 0.006 |
| 2D | Q RPE: O+ vs O-<br>SI RPE: O+ vs O- | 10 simulations | Paired t-test | $t_{(9)} = 1.19$<br>$t_{(9)} = 20.67$ | 0.26<br>$6.78 \times 10^{-9}$ |
| 2F | dLight: O+ vs O- | 41 sessions from<br>5 mice | Paired t-test | $t_{(66.4)} = 3.62$ | 0.000574 |
| 2G | $R^2$ of Q vs SI regression | 41 sessions from<br>5 mice | Paired t-test | $t_{(82)} = 2.2$ | 0.034 |
| 2J | Q RPE CPD vs<br>SI RPE CPD | 5 mice | Paired t-test | $t_{(4)} = 2.82$ | 0.047 |
| 2K | Q RPE alone $R^2$ vs<br>SI RPE alone $R^2$ | 5 mice | Paired t-test | $t_{(4)} = 4.34$ | 0.012 |
| 3B | p correct:<br>sham vs lesion | Sham: 25 sessions from<br>3 mice<br>Lesion: 35 sessions from<br>5 mice | t-test | $t_{(42)} = 3.55$ | 0.001 |
| 3C,D | Difference between mouse<br>and prediction from Q vs SI<br>Sham vs lesion | Sham: 25 sessions from<br>3 mice<br>Lesion: 35 sessions from<br>5 mice | Paired t-test<br>Paired t-test<br>Mixed effects ANOVA<br>Effect of Model<br>Effect of lesion<br>Interaction | $t_{(27)} = 7.21$<br>$t_{(54)} = 0.054$<br>$t_{(84)} = 0.065$<br>$t_{(84)} = 1.11$<br>$t_{(84)} = 2.49$ | $9.38 \times 10^{-8}$<br>0.96<br>0.95<br>0.27<br>0.01 |
| 3E | p SI consistent:<br>sham vs lesion | Sham: 25 sessions from<br>3 mice<br>Lesion: 35 sessions from<br>5 mice | t-test | $t_{(48)} = 3.75$ | 0.0005 |
| 3H | dLight: O+ vs O-<br>sham<br>lesion | 41 sessions from 5 mice<br>43 sessions from 5 mice | Paired t-test<br>Mixed effects ANOVA<br>Effect of past choice<br>Effect of lesion<br>Interaction | $t_{(66.4)} = 3.62$<br>$t_{(71.4)} = 0.65$<br>$t_{(137.6)} = 4.06$<br>$t_{(13.8)} = 0.82$<br>$t_{(137.8)} = 2.53$ | 0.000574<br>0.52<br>$8 \times 10^{-5}$<br>0.44<br>0.01 |
| 3I | R<sup>2</sup> of Q vs SI regression<br>sham<br>lesion | 41 sessions from 5 mice<br>43 sessions from 5 mice | Paired t-test | $t_{(60)} = 2.35$<br>$t_{(58.9)} = 0.30$ | 0.02<br>0.77 |
| 3J | Q RPE CPD vs<br>SI RPE CPD<br>sham<br>lesion | 5 mice<br>5 mice | Paired t-test<br>Mixed effects ANOVA<br>Effect of lesion<br>Effect of p context<br>Interaction | $t_{(4)} = 2.82$<br>$t_{(4)} = 2.00$<br>$F_{(1,8)} = 0.09$<br>$F_{(1,8)} = 0.62$<br>$F_{(1,8)} = 8.55$ | 0.047<br>0.12<br>0.77<br>0.45<br>0.01 |
| 4K | Decoding accuracy<br>vs chance<br>choice<br>value<br>context | 9 sessions from 3 mice | Paired t-test | $t_{(8)} = 5.33$<br>$t_{(8)} = 6.65$<br>$t_{(8)} = 6.13$ | 0.0007<br>0.0002<br>0.0003 |
| 4M | Decoding accuracy<br>vs chance<br>shuffle<br>test | 4 sessions from 2 mice<br>(ses. with > 100 neurons) | Paired t-test | $t_{(3)} = 1.06$<br>$t_{(3)} = 3.49$ | 0.37<br>0.03 |
| S1D | p incongruent trials<br>x switch | 10 mice | Repeated-measures ANOVA | $F_{(14,126)} = 7.98$ | $6.1 \times 10^{-12}$ |
| S1E | p incongruent choice<br>consistent with Q vs SI | 10 mice | Paired t-test | $t_{(9)} = 5.83$ | 0.0003 |
| S3B | p SI consistent after switch:<br>sham vs lesion | Sham: 25 sessions from<br>3 mice<br>Lesion: 35 sessions from<br>5 mice | t-test | $t_{(42)} = 2.44$ | 0.019 |
| S3B | p Q consistent after switch:<br>sham vs lesion | Sham: 25 sessions from<br>3 mice<br>Lesion: 35 sessions from<br>5 mice | t-test | $t_{(42)} = 1.31$ | 0.20 |
| S3C | p SI on incongruent choice:<br>sham vs lesion | Sham: 25 sessions from<br>3 mice<br>Lesion: 35 sessions from<br>5 mice | t-test | $t_{(8.5)} = 3.52$ | 0.007 |
| S3D | Parameters of model fits<br>sham vs lesion:<br>$\gamma$<br>c<br>bias | Sham: 25 sessions from<br>3 mice<br>Lesion: 35 sessions from<br>5 mice | t-test | $t_{(4)} = 5.47$<br>$t_{(4)} = 0.40$<br>$t_{(4)} = 0.51$ | 0.005<br>0.71<br>0.64 |
| S3E | p correct sham vs lesion | Sham: 41 sessions<br>from 5 mice<br>Lesion: 43 sessions<br>from 5 mice | t-test | $t_{(8)} = 0.36$ | 0.72 |
| S3F | Difference between mouse<br>and prediction from Q vs SI<br>Sham vs lesion | Sham: 41 sessions<br>from 5 mice<br>Lesion: 43 sessions<br>from 5 mice | t-test | $t_{(4)} = 26.32$<br>$t_{(4)} = 31.93$ | 0.00001<br>0.00001 |
| S3G | p SI consistent:<br>sham vs lesion | Sham: 41 sessions<br>from 5 mice<br>Caspase: 43 sessions<br>from 5 mice | t-test | $t_{(8)} = 0.20$ | 0.85 |

## REFERENCES

1. Wikenheiser, A.M., and Schoenbaum, G. (2016). Over the river, through the woods: cognitive maps in the hippocampus and orbitofrontal cortex. Nat Rev Neurosci 17, 513–523. 10.1038/nrn.2016.56.

2. Gershman, S.J., and Uchida, N. (2019). Believing in dopamine. Nat Rev Neurosci 20, 703–714. 10.1038/s41583-019-0220-7.

3. Wilson, R.C., Takahashi, Y.K., Schoenbaum, G., and Niv, Y. (2014). Orbitofrontal Cortex as a Cognitive Map of Task Space. Neuron 81, 267–279. 10.1016/j.neuron.2013.11.005.

4. Costa, V.D., Tran, V.L., Turchi, J., and Averbeck, B.B. (2015). Reversal Learning and Dopamine: A Bayesian Perspective. J Neurosci 35, 2407–2416. 10.1523/jneurosci.1989-14.2015.

5. Eckstein, M.K., Master, S.L., Dahl, R.E., Wilbrecht, L., and Collins, A.G.E. (2022). Reinforcement learning and Bayesian inference provide complementary models for the unique advantage of adolescents in stochastic reversal. Dev. Cogn. Neurosci. 55, 101106. 10.1016/j.dcn.2022.101106.

6. Babayan, B.M., Uchida, N., and Gershman, Samuel.J. (2018). Belief state representation in the dopamine system. Nat Commun 9, 1891. 10.1038/s41467-018-04397-0.

7. Starkweather, C.K., Babayan, B.M., Uchida, N., and Gershman, S.J. (2017). Dopamine reward prediction errors reflect hidden-state inference across time. Nat Neurosci 20, 581–589. 10.1038/nn.4520.

8. Vertechi, P., Lottem, E., Sarra, D., Godinho, B., Treves, I., Quendera, T., Lohuis, M.N.O., and Mainen, Z.F. (2020). Inference-Based Decisions in a Hidden State Foraging Task: Differential Contributions of Prefrontal Cortical Areas. Neuron 106, 166–176.e6. 10.1016/j.neuron.2020.01.017.

9. Qü, A.J., Tai, L.-H., Hall, C.D., Tu, E.M., Eckstein, M.K., Mishchanchuk, K., Lin, W.C., Chase, J.B., MacAskill, A.F., Collins, A.G.E., et al. (2023). Nucleus accumbens dopamine release reflects Bayesian inference during instrumental learning. bioRxiv, 2023.11.10.566306. 10.1101/2023.11.10.566306.

10. Lak, A., Nomoto, K., Keramati, M., Sakagami, M., and Kepecs, A. (2017). Midbrain Dopamine Neurons Signal Belief in Choice Accuracy during a Perceptual Decision. Curr Biol 27, 821–832. 10.1016/j.cub.2017.02.026.

11. Adams, R.A., Huys, Q.J.M., and Roiser, J.P. (2015). Computational Psychiatry: towards a mathematically informed understanding of mental illness. J Neurology Neurosurg Psychiatry 87, 53. 10.1136/jnnp-2015-310737.

12. Schlagenhauf, F., Huys, Q.J.M., Deserno, L., Rapp, M.A., Beck, A., Heinze, H.-J., Dolan, R., and Heinz, A. (2014). Striatal dysfunction during reversal learning in unmedicated schizophrenia patients. Neuroimage 89, 171–180. 10.1016/j.neuroimage.2013.11.034.

13. Huys, Q.J., Pizzagalli, D.A., Bogdan, R., and Dayan, P. (2013). Mapping anhedonia onto reinforcement learning: a behavioural meta-analysis. Biology Mood Anxiety Disord 3, 12. 10.1186/2045-5380-3-12.

14. Mkrtchian, A., Aylward, J., Dayan, P., Roiser, J.P., and Robinson, O.J. (2017). Modeling Avoidance in Mood and Anxiety Disorders Using Reinforcement Learning. Biol Psychiat 82, 532–539. 10.1016/j.biopsych.2017.01.017.

15. Radulescu, A., and Niv, Y. (2019). State representation in mental illness. Curr Opin Neurobiol 55, 160–166. 10.1016/j.conb.2019.03.011.

16. O’Keefe, J., and Nadel, L. (1978). The Hippocampus as a Cognitive Map (Oxford University Press.).

17. Hartley, T., Lever, C., Burgess, N., and O’Keefe, J. (2014). Space in the brain: how the hippocampal formation supports spatial cognition. Philosophical Transactions Royal Soc B Biological Sci 369, 20120510. 10.1098/rstb.2012.0510.

18. Per, A., Richard, M., David, A., Tim, B., and John, O. (2006). The Hippocampus Book 10.1093/acprof:oso/9780195100273.003.0002.

19. Sanders, H., Wilson, M.A., and Gershman, S.J. (2020). Hippocampal remapping as hidden state inference. Elife 9, e51140. 10.7554/elife.51140.

20. Courellis, H.S., Mixha, J., Cardenas, A.R., Kimmel, D., Reed, C.M., Valiante, T.A., Salzman, C.D., Mamelak, A.N., Fusi, S., and Rutishauser, U. (2023). Abstract representations emerge in human hippocampal neurons during inference behavior. bioRxiv, 2023.11.10.566490. 10.1101/2023.11.10.566490.

21. Tai, L.-H., Lee, A.M., Benavidez, N., Bonci, A., and Wilbrecht, L. (2012). Transient stimulation of distinct subpopulations of striatal neurons mimics changes in action value. Nat Neurosci 15, 1281–1289. 10.1038/nn.3188.

22. Parker, N.F., Baidya, A., Cox, J., Haetzel, L.M., Zhukovskaya, A., Murugan, M., Engelhard, B., Goldman, M.S., and Witten, I.B. (2022). Choice-selective sequences dominate in cortical relative to thalamic inputs to NAc to support reinforcement learning. Cell Reports 39, 110756. 10.1016/j.celrep.2022.110756.

23. Parker, N.F., Cameron, C.M., Taliaferro, J.P., Lee, J., Choi, J.Y., Davidson, T.J., Daw, N.D., and Witten, I.B. (2016). Reward and choice encoding in terminals of midbrain dopamine neurons depends on striatal target. Nat Neurosci 19, 845–854. 10.1038/nn.4287.

24. Stephenson-Jones, M., Yu, K., Ahrens, S., Tucciarone, J.M., Huijstee, A.N. van, Mejia, L.A., Penzo, M.A., Tai, L.-H., Wilbrecht, L., and Li, B. (2016). A basal ganglia circuit for evaluating action outcomes. Nature 539, 289–293. 10.1038/nature19845.

25. Hattori, R., Danskin, B., Babic, Z., Mlynaryk, N., and Komiyama, T. (2019). Area-Specificity and Plasticity of History-Dependent Value Coding During Learning. Cell 177, 1858–1872.e15. 10.1016/j.cell.2019.04.027.

26. Miller, K.J., Botvinick, M.M., and Brody, C.D. (2021). From predictive models to cognitive models: Separable behavioral processes underlying reward learning in the rat. bioRxiv, 461129. 10.1101/461129.

27. Beron, C.C., Neufeld, S.Q., Linderman, S.W., and Sabatini, B.L. (2022). Mice exhibit stochastic and efficient action switching during probabilistic decision making. Proc. Natl. Acad. Sci. 119, e2113961119. 10.1073/pnas.2113961119.

28. Groman, S.M., Keistler, C., Keip, A.J., Hammarlund, E., DiLeone, R.J., Pittenger, C., Lee, D., and Taylor, J.R. (2019). Orbitofrontal Circuits Control Multiple Reinforcement-Learning Processes. Neuron 103, 734-746.e3. 10.1016/j.neuron.2019.05.042.

29. Lak, A., Okun, M., Moss, M.M., Gurnani, H., Farrell, K., Wells, M.J., Reddy, C.B., Kepecs, A., Harris, K.D., and Carandini, M. (2020). Dopaminergic and Prefrontal Basis of Learning from Sensory Confidence and Reward Value. Neuron 105, 700–711.e6. 10.1016/j.neuron.2019.11.018.

30. Sadacca, B.F., Jones, J.L., and Schoenbaum, G. (2016). Midbrain dopamine neurons compute inferred and cached value prediction errors in a common framework. Elife 5, e13665. 10.7554/elife.13665.

31. Patriarchi, T., Cho, J.R., Merten, K., Howe, M.W., Marley, A., Xiong, W.-H., Folk, R.W., Broussard, G.J., Liang, R., Jang, M.J., et al. (2018). Ultrafast neuronal imaging of dopamine dynamics with designed genetically encoded sensors. Science 360, eaat4422. 10.1126/science.aat4422.

32. Komorowski, R.W., Garcia, C.G., Wilson, A., Hattori, S., Howard, M.W., and Eichenbaum, H. (2013). Ventral Hippocampal Neurons Are Shaped by Experience to Represent Behaviorally Relevant Contexts. J Neurosci 33, 8079–8087. 10.1523/jneurosci.5458-12.2013.

33. Behrens, T.E.J., Muller, T.H., Whittington, J.C.R., Mark, S., Baram, A.B., Stachenfeld, K.L., and Kurth-Nelson, Z. (2018). What Is a Cognitive Map? Organizing Knowledge for Flexible Behavior. Neuron 100, 490–509. 10.1016/j.neuron.2018.10.002.

34. Wee, R.W.S., and MacAskill, A.F. (2020). Biased Connectivity of Brain-wide Inputs to Ventral Subiculum Output Neurons. Cell Reports 30, 3644–3654.e6. 10.1016/j.celrep.2020.02.093.

35. Turner, V.S., O’Sullivan, R.O., and Kheirbek, M.A. (2022). Linking external stimuli with internal drives: A role for the ventral hippocampus. Curr Opin Neurobiol 76, 102590. 10.1016/j.conb.2022.102590.

36. Duvelle, É., Grieves, R.M., and Meer, M.A. van der (2023). Temporal context and latent state inference in the hippocampal splitter signal. eLife 12, e82357. 10.7554/elife.82357.

37. Gershman, S.J., Blei, D.M., and Niv, Y. (2010). Context, Learning, and Extinction. Psychol Rev 117, 197–209. 10.1037/a0017808.

38. Kubie, J.L., Levy, E.R.J., and Fenton, A.A. (2020). Is hippocampal remapping the physiological basis for context? Hippocampus 30, 851–864. 10.1002/hipo.23160.

39. Fuhs, M.C., and Touretzky, D.S. (2007). Context Learning in the Rodent Hippocampus. Neural Comput 19, 3173–3215. 10.1162/neco.2007.19.12.3173.

40. Sharpe, M.J., Wikenheiser, A.M., Niv, Y., and Schoenbaum, G. (2015). The State of the Orbitofrontal Cortex. Neuron 88, 1075–1077. 10.1016/j.neuron.2015.12.004.

41. Zhou, J., Jia, C., Montesinos-Cartagena, M., Gardner, M.P.H., Zong, W., and Schoenbaum, G. (2021). Evolving schema representations in orbitofrontal ensembles during learning. Nature 590, 606–611. 10.1038/s41586-020-03061-2.

42. Sadacca, B.F., Wied, H.M., Lopatina, N., Saini, G.K., Nemirovsky, D., and Schoenbaum, G. (2018). Orbitofrontal neurons signal sensory associations underlying model-based inference in a sensory preconditioning task. Elife 7, e30373. 10.7554/elife.30373.

43. Starkweather, C.K., Gershman, S.J., and Uchida, N. (2018). The Medial Prefrontal Cortex Shapes Dopamine Reward Prediction Errors under State Uncertainty. Neuron 98, 616–629.e6. 10.1016/j.neuron.2018.03.036.

44. Wikenheiser, A.M., Marrero-Garcia, Y., and Schoenbaum, G. (2017). Suppression of Ventral Hippocampal Output Impairs Integrated Orbitofrontal Encoding of Task Structure. Neuron 95, 1197–1207.e3. 10.1016/j.neuron.2017.08.003.

45. Sánchez-Bellot, C., AlSubaie, R., Mishchanchuk, K., Wee, R.W.S., and MacAskill, A.F. (2022). Two opposing hippocampus to prefrontal cortex pathways for the control of approach and avoidance behaviour. Nat. Commun. 13, 339. 10.1038/s41467-022-27977-7.

46. AlSubaie, R., Wee, R.W., Ritoux, A., Mishchanchuk, K., Passlack, J., Regester, D., and MacAskill, A.F. (2021). Control of parallel hippocampal output pathways by amygdalar long-range inhibition. Elife 10, e74758. 10.7554/elife.74758.

47. Wee, R.W.S., Mishchanchuk, K., AlSubaie, R., Church, T.W., Gold, M.G., and MacAskill, A.F. (2024). Internal-state-dependent control of feeding behavior via hippocampal ghrelin signaling. Neuron 112, 288–305.e7. 10.1016/j.neuron.2023.10.016.

48. LeGates, T.A., Kvarta, M.D., Tooley, J.R., Francis, T.C., Lobo, M.K., Creed, M.C., and Thompson, S.M. (2018). Reward behaviour is regulated by the strength of hippocampus–nucleus accumbens synapses. Nature 564, 258–262. 10.1038/s41586-018-0740-8.

49. Ciocchi, S., Passecker, J., Malagon-Vina, H., Mikus, N., and Klausberger, T. (2015). Selective information routing by ventral hippocampal CA1 projection neurons. Science 348, 560–563. 10.1126/science.aaa3245.

50. Floresco, S.B., Todd, C.L., and Grace, A.A. (2001). Glutamatergic afferents from the hippocampus to the nucleus accumbens regulate activity of ventral tegmental area dopamine neurons. The Journal of neuroscience : the official journal of the Society for Neuroscience 21, 4915–4922.

51. Trouche, S., Koren, V., Doig, N.M., Ellender, T.J., El-Gaby, M., Lopes-dos-Santos, V., Reeve, H.M., Perestenko, P.V., Garas, F.N., Magill, P.J., et al. (2019). A Hippocampus-Accumbens Tripartite Neuronal Motif Guides Appetitive Memory in Space. Cell 176, 1393–1406.e16. 10.1016/j.cell.2018.12.037.

52. Nour, M.M., Liu, Y., Arumuham, A., Kurth-Nelson, Z., and Dolan, R.J. (2021). Impaired neural replay of inferred relationships in schizophrenia. Cell 184, 4315–4328.e17. 10.1016/j.cell.2021.06.012.

53. Godsil, B.P., Kiss, J.P., Spedding, M., and Jay, T.M. (2013). The hippocampal–prefrontal pathway: The weak link in psychiatric disorders? Eur Neuropsychopharm 23, 1165–1181. 10.1016/j.euroneuro.2012.10.018.

54. Campbell, S., and Macqueen, G. (2004). The role of the hippocampus in the pathophysiology of major depression. J. psychiatry Neurosci. : JPN 29, 417–426.

55. Mukherjee, A., Carvalho, F., Eliez, S., and Caroni, P. (2019). Long-Lasting Rescue of Network and Cognitive Dysfunction in a Genetic Schizophrenia Model. Cell 178, 1387–1402.e14. 10.1016/j.cell.2019.07.023.

56. Sigurdsson, T., Stark, K.L., Karayiorgou, M., Gogos, J.A., and Gordon, J.A. (2010). Impaired hippocampal–prefrontal synchrony in a genetic mouse model of schizophrenia. Nature 464, 763–767. 10.1038/nature08855.

57. Husain, M., and Roiser, J.P. (2018). Neuroscience of apathy and anhedonia: a transdiagnostic approach. Nat Rev Neurosci 19, 470–484. 10.1038/s41583-018-0029-9.

58. Resendez, S.L., Jennings, J.H., Ung, R.L., Namboodiri, V.M.K., Zhou, Z.C., Otis, J.M., Nomura, H., McHenry, J.A., Kosyk, O., and Stuber, G.D. (2016). Visualization of cortical, subcortical and deep brain neural circuit dynamics during naturalistic mammalian behavior with head-mounted microscopes and chronically implanted lenses. Nat Protoc 11, 566–597. 10.1038/nprot.2016.021.

59. Lau, B., and Glimcher, P.W. (2005). DYNAMIC RESPONSE-BY-RESPONSE MODELS OF MATCHING BEHAVIOR IN RHESUS MONKEYS. J Exp Anal Behav 84, 555–579. 10.1901/jeab.2005.110-04.

60. Wilson, R.C., and Collins, A.G. (2019). Ten simple rules for the computational modeling of behavioral data. Elife 8, e49547. 10.7554/elife.49547.

61. Jeong, Y., Huh, N., Lee, J., Yun, I., Lee, J.W., Lee, I., and Jung, M.W. (2018). Role of the hippocampal CA1 region in incremental value learning. Sci Rep-uk 8, 9870. 10.1038/s41598-018-28176-5.

62. Li, J., Schiller, D., Schoenbaum, G., Phelps, E.A., and Daw, N.D. (2011). Differential roles of human striatum and amygdala in associative learning. Nat Neurosci 14, 1250–1252. 10.1038/nn.2904.

63. Costa, V.D., Dal Monte, O., Lucas, D.R., Murray, E.A., and Averbeck, B.B. (2016). Amygdala and Ventral Striatum Make Distinct Contributions to Reinforcement Learning. Neuron 92, 505–517. 10.1016/j.neuron.2016.09.025.

64. Blanco-Pozo, M., Akam, T., and Walton, M.E. (2023). Dopamine-independent state inference mediates expert reward guided decision making. bioRxiv, 2021.06.25.449995. 10.1101/2021.06.25.449995.

65. Dong, Z., Mau, W., Feng, Y., Pennington, Z.T., Chen, L., Zaki, Y., Rajan, K., Shuman, T., Aharoni, D., and Cai, D.J. (2022). Minian, an open-source miniscope analysis pipeline. Elife 11, e70661. 10.7554/elife.70661.

66. Zhou, P., Resendez, S.L., Rodriguez-Romaguera, J., Jimenez, J.C., Neufeld, S.Q., Giovannucci, A., Friedrich, J., Pnevmatikakis, E.A., Stuber, G.D., Hen, R., et al. (2018). Efficient and accurate extraction of in vivo calcium signals from microendoscopic video data. eLife 7, e28728. 10.7554/elife.28728.

67. Ahmed, M.S., Priestley, J.B., Castro, A., Stefanini, F., Canales, A.S.S., Balough, E.M., Lavoie, E., Mazzucato, L., Fusi, S., and Losonczy, A. (2020). Hippocampal Network Reorganization Underlies the Formation of a Temporal Association Memory. Neuron 107, 283–291.e6. 10.1016/j.neuron.2020.04.013.

68. Akam, T., Rodrigues-Vaz, I., Marcelo, I., Zhang, X., Pereira, M., Oliveira, R.F., Dayan, P., and Costa, R.M. (2020). The Anterior Cingulate Cortex Predicts Future States to Mediate Model-Based Action Selection. Neuron. 10.1016/j.neuron.2020.10.013.

69. Bernardi, S., Benna, M.K., Rigotti, M., Munuera, J., Fusi, S., and Salzman, C.D. (2020). The Geometry of Abstraction in the Hippocampus and Prefrontal Cortex. Cell 183, 954–967.e21. 10.1016/j.cell.2020.09.031.

